# Directed evolution improves the catalytic efficiency of TEV protease

**DOI:** 10.1101/811570

**Authors:** Mateo I Sanchez, Alice Y Ting

**Author notes:** Correspondence should be addressed to A. Y. T.

## Abstract

Tobacco etch virus protease (TEV) is one of the most widely-used proteases in biotechnology because of its exquisite sequence-specificity. A limitation, however, is its slow catalytic rate. We developed a generalizable yeast-based platform for directed evolution of protease catalytic properties. Protease activity is read out via proteolytic release of a membrane-anchored transcription factor, and we temporally regulate access to TEV’s cleavage substrate using a photosensory LOV domain. By gradually decreasing light exposure time, we enriched faster variants of TEV over multiple rounds of selection. Our S153N mutant (uTEV1Δ), when incorporated into the calcium integrator FLARE, improved the signal/background ratio by 27-fold, and enabled recording of neuronal activity in culture with 60-second temporal resolution. Given the widespread use of TEV in biotechnology, both our evolved TEV mutants and the directed evolution platform used to generate them, could be beneficial across a wide range of applications.

## Main text

Proteases are ubiquitous in biology, frequently initiating or terminating endogenous signaling cascades. Their peptide bond cleavage activities have been harnessed for a wide range of biotechnological applications, including bottom-up mass spectrometry (MS)-based proteomics, affinity purification, neuronal silencing (e.g., botulinum protease [1]), light-regulated apoptosis [2], tagging of newly synthesized proteins [3], assembly of protein droplets [4], protease-based synthetic circuits [5,6], regulation of TALENs [7], transcriptional readout of calcium [8,9] and protein-protein interactions [10,11,12].

One of the most frequently-used proteases in biotechnology is TEV, the 27 kD cysteine protease from tobacco etch virus. TEV is appealing because it is active in the mammalian cytosol, has no required cofactors, recognizes a 7-amino acid consensus peptide substrate, and, most importantly, is highly sequence-specific, exhibiting negligible activity towards endogenous mammalian proteomes. Consequently, TEV has been harnessed for sequence-specific transcription factor release in response to calcium and light in FLARE [8], GPCR activation in TANGO [10], and GPCR activation and light in SPARK [11, 12]. In the recently reported CHOMP [5] and SPOC [6] tools, TEV is activated by inputs such as rapamycin or abscisic acid.

Despite the exquisite sequence-specificity of TEV, a major limitation is its slow catalysis. With a *k*_*cat*_ of 0.18 s^−1^ (for its best high-affinity substrate sequence, ENLYFQS [13]), TEV is slower than many other proteases used for biotechnology, such as trypsin (*k*_*cat*_ 75 s^−1^ [14]) and subtilisin (*k*_*cat*_ 50 s^−1^ [15]). This slow turnover fundamentally limits the performance of technologies that rely on TEV, such as FLARE [8]. In vivo, FLARE requires calcium and light stimulation for at least 30 minutes to give TEV sufficient time to release detectable quantities of membrane-anchored transcription factor [8]. Yet for the neuronal activity integration applications for which FLARE is designed, a temporal resolution of just a few minutes, or even seconds, is desired – a goal we have found impossible to achieve using wild-type TEV (vide infra).

There have not been systematic efforts to improve the catalytic rate of TEV, apart from optimization of its substrate sequence (TEVcs). Directed evolution has previously been applied to alter TEV’s sequence *specificity*, producing variants that have either similar [16] or depressed [17] catalytic efficiency compared to wild-type TEV. Here, the goal of our work is a generalizable strategy for improving the catalytic efficiency of proteases of biotechnological interest via directed evolution. After applying our platform across multiple rounds of selection to two TEV mutant libraries, we enriched three proteases, named uTEV1Δ, uTEV2Δ, and uTEV3, that have improved catalytic efficiency for TEVcs cleavage compared to wild type TEV. uTEV1Δ was evaluated in the context of FLARE and SPARK, and shown to improve the performance of these tools in mammalian cells.

## Results

### A yeast-based platform for evolving protease catalysis

Yeast are attractive as a platform for directed evolution because they naturally compartmentalize chemical reactions and can be sorted by FACS instruments over a large dynamic range. We previously used yeast-based directed evolution to improve the properties of APEX peroxidase [18], promiscuous biotin ligase [19], split horseradish peroxidase [20], and split APEX [21]. Iverson et al. developed a yeast platform to alter the sequence-specificity of TEV [17]. In their approach, a TEV mutant library was co-expressed in the yeast ER lumen with a HA-tagged reporter linked by a TEVcs sequence to an ER retention motif. An active TEV mutant could remove the ER retention motif and allow the reporter to traffic to the cell surface, where it could be detected by a fluorescent anti-His_6_ antibody. While this scheme was effective for discovering TEV variants with activity towards altered TEVcs sequences, it was not able to enrich highly active proteases over moderately active ones. This is because the time window for TEV action on TEVcs was not controlled; TEV mutants could act on TEVcs over >8 hours (the time window for co-expression), enabling even low-activity mutants to be enriched.

To devise a platform that could be used to enrich faster proteases over moderately-active ones, we implemented the following: (1) we moved the system into the yeast cytosol, since this reducing environment more closely resembles the eventual context in which evolved TEVs will mostly be used, (2) we coupled TEV activity to the release of a membrane-anchored transcription factor (TF) which in turn drives the expression of a fluorescent protein reporter, in order to increase sensitivity and dynamic range of the protease activity readout, (3) we fused TEV and TEVcs to the photoinducible protein binding pair CRY-CIBN [22] so that our selections can be applied to truncated, low-affinity versions of TEV that are utilized in FLARE and SPARK tools; despite the low affinity, recognition of TEVcs by TEV can be induced by blue light activation of CRY-CIBN. And most importantly, (4), we photocaged the TEVcs substrate sequence with an improved LOV domain (eLOV [8]) in order to exert control over the time window TEV is available to cleave TEVcs.

The scheme in Figure 1A shows the design of our protease evolution platform in yeast. The TEV mutant library is expressed as a fusion to CRY and mCherry. The transcription factor (TF) used to read out TEV activity is linked to a plasma membrane anchor via CIBN and LOV-caged TEVcs. Upon irradiation of the cell population with blue light, CRY and CIBN form an intermolecular complex, driving TEV into proximity of TEVcs. In addition, the LOV domain undergoes a conformational change, exposing TEVcs. Active TEV mutants will then cleave TEVcs, releasing the TF for translocation to the nucleus and transcription of the reporter gene (Citrine). Six hours later (to allow time for Citrine transcription, translation, and maturation) FACS is used to enrich yeast cells with high Citrine/mCherry intensity ratio, which is indicative of high TEV cleavage activity (Figure 1B). The central feature of our platform is that selection stringency can be increased simply by decreasing the blue light irradiation time. With less time available to sterically access TEVcs, only the most active TEV mutants will produce Citrine signal over background.

**Figure 1:**
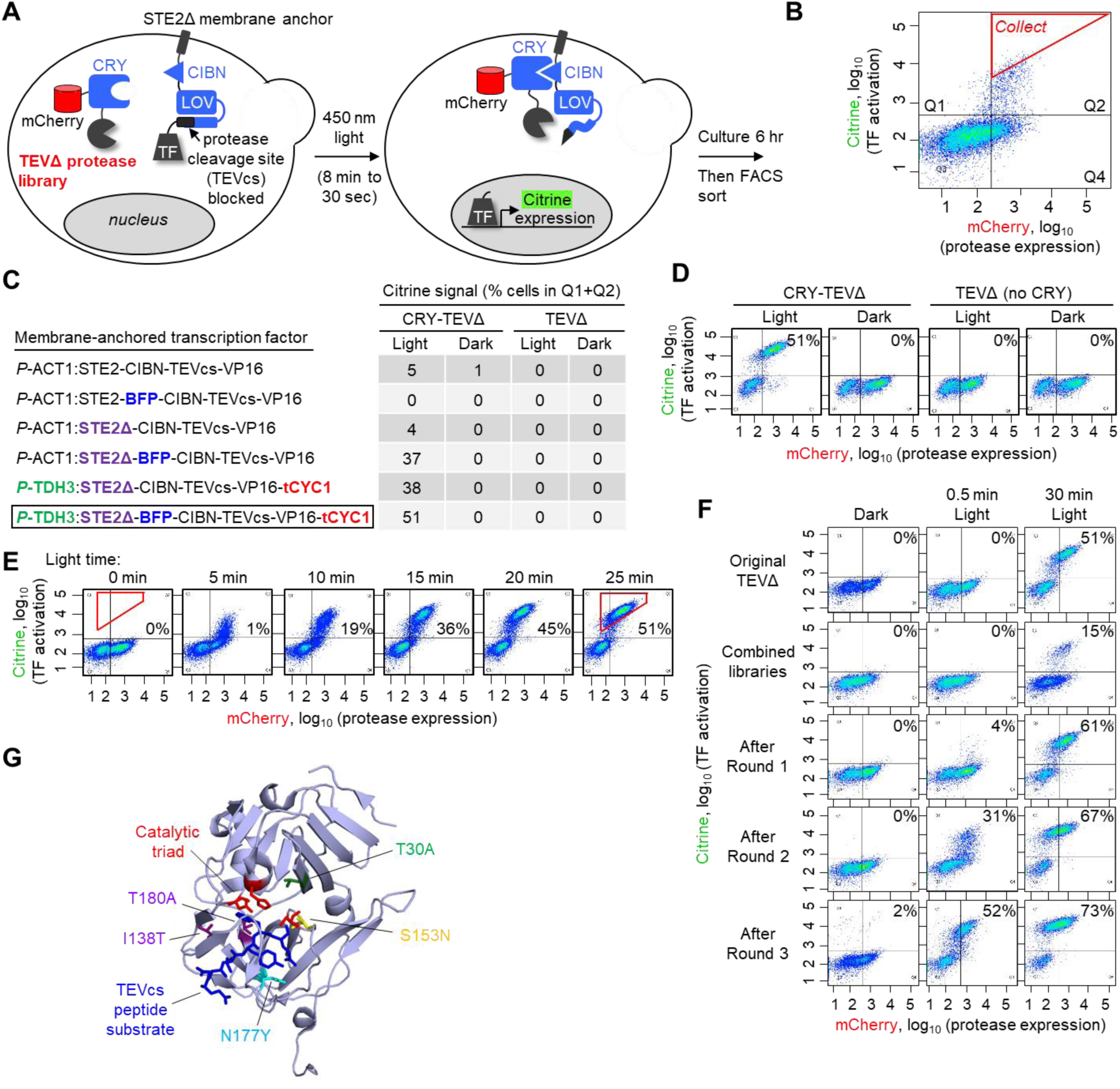
Yeast platform for directed evolution of high-turnover, low-affinity proteases A. Schematic of evolution platform in the yeast cytosol. A library of truncated TEV protease (TEVΔ) variants is fused to CRY and mCherry. A transcription factor (TF) is tethered to the plasma membrane via a TEV cleavage site (TEVcs), a LOV domain, and CIBN. Upon exposure of cells to blue 450 nm light, the CRY-CIBN interaction brings the TEV protease into proximity of TEVcs, and the LOV domain changes conformation to expose TEVcs. Proteolysis releases the TF, which translocates to the nucleus and drives expression of the reporter gene Citrine. Selection stringency can be increased by reducing the irradiation time (allowing less time for TEV-catalyzed TF release). B. FACS analysis of yeast library 6 hours after 8-minute blue light exposure. A subpopulation of cells display Citrine fluorescence above background, indicating that they contain active TEV. mCherry is used to read out protease expression levels. The red gate was used to collect cells with the highest Citrine/mCherry intensity ratios. C. Optimization of membrane-anchored transcription factor component of the evolution platform. For each construct, FACS analysis was performed as shown in (B), 6 hours after 45-minute blue light exposure. D. Controls are shown with light omitted (columns 2 and 4) or CRY omitted (columns 3-4). Table values reflect the fraction of cells with high Citrine intensity, i.e., cells in the upper FACS quadrants Q1 and Q2 (quadrants are defined in (B)). D. FACS plots corresponding to the last row of the table in (C). All other FACS plots are shown in **Supplementary Figure 1B**. This experiment was performed twice with similar results. E. Citrine signal scales with light irradiation time. As the 450 nm light exposure time is increased from 0 min to 25 min, the resulting Citrine expression 6 hours later increases. Values in each plot reflect the percentage of cells within the red polygonal gate shown. This experiment was performed twice with similar results. F. FACS plots summarizing the progress of the selections. Re-amplified yeast pools were analyzed side by side under the three conditions shown (three columns). Values reflect the fraction of Citrine-positive cells, i.e. cells in upper quadrants Q1 and Q2. Additional FACS plots and summary graph in **Supplementary Figure 2A**. This experiment was performed once. G. Mutations enriched by the evolution, highlighted on a ribbon structure of wild-type TEV protease (PDB: 1LVM [24]) in complex with its peptide substrate (in dark blue). uTEV1Δ contains the mutation S153N, while uTEV2Δ has both S153N and T30A mutations. From our high-affinity TEV evolution (Figure 3), we also enriched the mutations I138T and T180A. (uTEV3 has three mutations: I138T, S153N, and T180A). The N177Y mutation is also highlighted because it arose during selections, although we rejected it because it increases affinity for TEVcs.

Using C-terminally truncated, low-affinity wild-type TEV (TEVΔ219 [23], or “TEVΔ”) as our starting template, we optimized a number of features of the platform (see Supplementary Text 1 and Figures 1C-D). After optimization, we observed that Citrine reporter signal increased as the blue light irradiation time was lengthened from 5 to 30 minutes (Figure 1E), suggesting that the selection stringency of this platform is tunable by modulating the light time.

To implement the directed evolution, we first generated a library of TEVΔ mutants using error-prone PCR. Sequencing indicated an average mutation rate of 4 amino acids per gene. The library was transformed into yeast cells along with a membrane-anchored TF bearing a low-affinity TEVcs sequence (ENLYFQ/M), and we performed three successive rounds of selection. The blue light time was gradually decreased from 8 minutes to 30 seconds, and cells with high Citrine/mCherry signal ratio were enriched by FACS. Yeast populations collected after each round were amplified and compared under matched conditions. Figure 1F shows that the post-round 3 population is much more active than both the original library and wild-type TEVΔ.

### Characterization of evolved TEVΔ mutants

Sequencing after round 3 showed that specific mutations were enriched by the selection. Excitingly, several of these mutations (T30A, T30I, S31W and S153N) surround the catalytic triad (Figure 1G) in the wild-type TEV structure (PDB: 1LVM) [24], and might reasonably be expected to lower the energy of the transition state, improving catalysis (Figure 1G). An additional enriched mutation, N117Y, interacts directly with the bound TEVcs. We hypothesize that N117Y was enriched because it enhances binding to TEVcs instead of improving k_cat_ (also supported by data below in Figure 2A). This outcome is undesirable in our pursuit of a fast-turnover but low-affinity (proximity-dependent) TEV variant.

**Figure 2:**
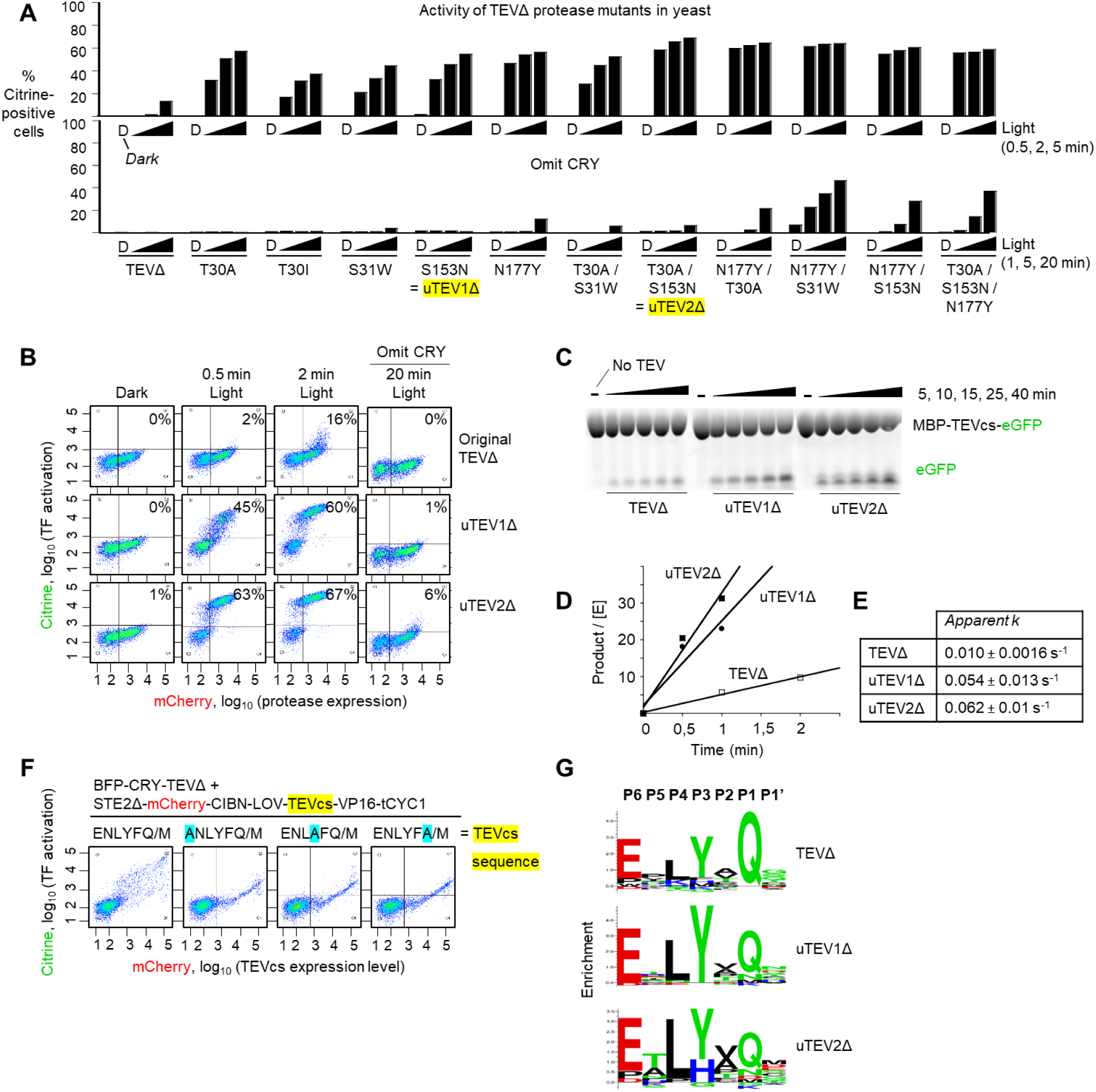
Characterization of evolved low-affinity proteases (uTEV1Δ and uTEV2Δ) in yeast and in vitro. A Comparison of evolved single, double, and triple TEVΔ mutants in yeast, with CRY present (top) or omitted (bottom; to test proximity-dependence of cleavage). Experiment was performed as in Figure 1A and FACS plots were quantified as in Figure 1C. For each clone, three irradiation times were tested (0.5, 2, and 5 min) in addition to the dark state (D). The two clones with the highest proximity-dependent activity are highlighted yellow. Additional timepoints and FACS plots shown in **Supplementary Figure 3**. B FACS plots for the two best clones in (A). Percentages show the fraction of Citrine-positive cells in Q1 + Q2. C Fluorescent gel measuring the kinetics of purified TEV proteases. The substrate protein MBP-TEVcs-GFP was incubated with the indicated TEV mutants (MBP = maltose binding protein; TEVcs = ENLYFQ/M). At various timepoints, the reaction was quenched, run on SDS-PAGE, and visualized by in-gel fluorescence. [MBP-TEVcs-GFP] was 360 μM and all proteases were at 750 nM. This experiment was performed independently three times with similar results. D Quantitation of protease reaction rates, using the fluorescent gel assay in (C). E Apparent rate constants based on initial velocity measurements in (D). Due to protein solubility limits, the maximum concentration of TEVcs was 360 μM (much lower than the expected K_m_). Therefore, the values likely represent lower bounds to the actual k_cat_. Three technical replicates were performed per condition. F Profiling protease sequence-specificity in yeast. Setup was the same as in Figure 1A, except the TEVcs sequence is randomized, and mCherry is fused to TEVcs rather than to TEV in order to quantify TEVcs expression level (schematic in **Supplementary Figure 5A**). The FACS plots show the cleavage extent for various TEVcs test substrates, 6 hours after 30-minute blue light irradiation. Forward slash indicates proteolysis site. Mutations at the −6, −3, and −1 positions of TEVcs greatly reduce cleavage by wild-type TEVΔ. This experiment was performed once. G Sequence specificity profiles of wild-type TEVΔ, uTEV1Δ, and uTEV2Δ obtained via sequencing of FACS-enriched TEVcs libraries (seven TEVcs libraries for each protease variant). FACS plots and sequencing data in **Supplementary Figure 5C**.

To evaluate the activities of evolved TEV mutants, we first used our yeast setup. Figure 2A shows that all our evolved mutants are significantly more active than wild-type TEVΔ, producing greater Citrine expression at all time points. To check for interaction-dependence of the cleavage activity, we repeated the assay with TEV mutants fused to mCherry only and not mCherry-CRY (“omit CRY” control, Figure 2A). Whereas most of these lost Citrine signal, the mutants containing the N177Y mutation still exhibited activity. The ability of these N177Y mutants (which are all truncated at position 219) to cleave TEVcs even when the CRY-CIBN pair is not present to bring the protease and substrate together under blue light, is consistent with the notion that N177Y may increase the affinity of TEVΔ for TEVcs. From our analysis in yeast, the two TEV mutants with the highest *proximity-dependent* activity were S153N (“uTEV1Δ”) and the T30A/S153N double mutant (“uTEV2Δ”) (Figure 2B and Supplementary Figure 3).

**Figure 3:**
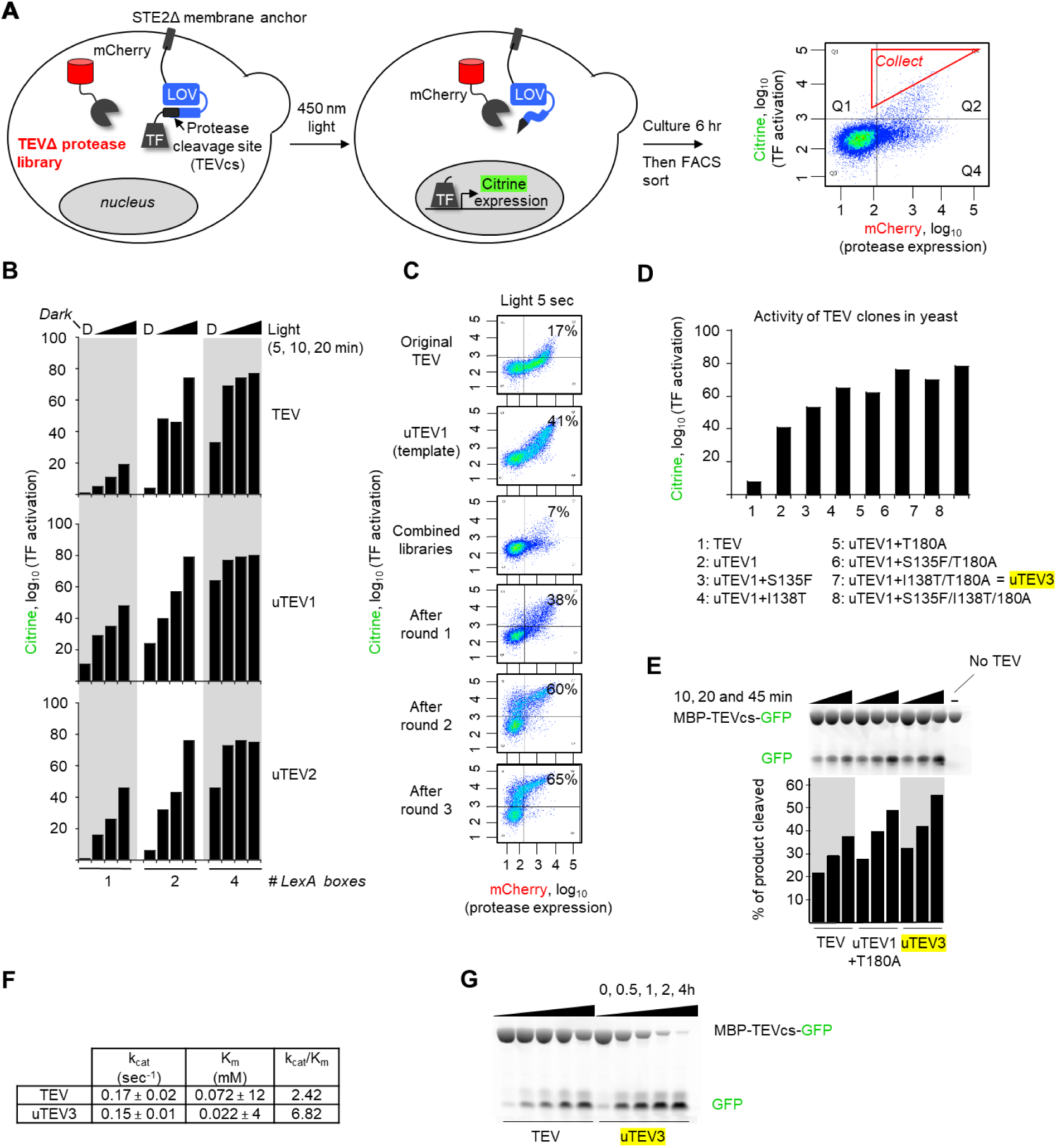
Yeast platform applied to the evolution of high-affinity proteases A. Selection scheme in yeast cytosol. A library of full-length TEV variants is expressed as a fusion to mCherry. The transcription factor (TF) is anchored to the plasma membrane via a protease-sensitive linker. FACS is used to enrich cells with high Citrine/mCherry intensity ratios. On the right is a FACS plot from the first round of evolution. The red gate shows the cells with high Citrine/mCherry intensity ratio that were collected by FACS. B. Tuning dynamic range of the evolution platform. By varying the number of LexA boxes in the promoter recognized by the LexA-VP16 TF, we modulated the sensitivity of the Citrine readout. More LexA boxes resulted in higher sensitivity, i.e., higher Citrine expression in response to short light exposure times. Corresponding FACS data is in **Supplementary Figures 6**. C. Results of selection. Selection was performed using the high-affinity TEVcs ENLYFQ/S. Percentages show fraction of Citrine-positive cells in Q1+Q2. Additional FACS plots and conditions are in **Supplementary Figure 7A**. D. Analysis of individual clones enriched by the selection. Activities were quantified in yeast by Citrine expression level, as in Figure 1F. Additional characterization in yeast in **Supplementary Figure 8**. E. Fluorescent gel assay measuring the kinetics of purified proteases. The protein substrate MBP-TEVcs-GFP (72 kDa, 28 μM, TEVcs = ENLYFQ/S) was incubated with the indicated proteases (all full-length, 125 nM) for 10, 20, or 45 min before SDS-PAGE and in-gel fluorescence detection of GFP. This experiment was performed once. F. Kinetic parameters for wild-type TEV and uTEV3 (containing the mutations I138T, S153N, and T180A), obtained via the fluorescence gel assay shown in (E). The MBP-TEVcs-GFP substrate concentration was varied from 7.5 to 320 μM to obtain the K_m_. Michaelis-Menten plots are in **Supplementary Figure 4**. Three technical replicates were performed per condition. G. uTEV3 is more efficient than wild-type TEV for affinity tag removal. MBP-TEVcs-GFP (72 kDa, 10 μM, TEVcs = ENLYFQ/S) was incubated with wild-type TEV or uTEV3 for 0-4 h. The product was analyzed by SDS-PAGE and in-gel fluorescence. This experiment was performed independently two times with similar results.

Next, we characterized our evolved TEV mutants in vitro. uTEV1Δ and uTEV2Δ, along with wild-type TEVΔ, were expressed and purified from bacteria (Supplementary Figure 4) and combined with the substrate protein MBP-TEVcs-GFP (MBP is maltose binding protein and TEVcs is the low-affinity substrate sequence ENLYFQ/M used in the yeast selection). The solubility of MBP-TEVcs-GFP limited its maximum concentration to 360 μM, which is likely far below the TEVΔ saturation concentration (for reference, wild-type TEVΔ has a K_m_ of 448 μM for the *high-affinity* TEVcs ENLYFQ/S [23]; its K_m_ for the low-affinity TEVcs used here has never been measured but is likely much higher). After incubation for various times, we evaluated the cleavage extent by running the reactions on SDS-PAGE and performing in-gel fluorescence imaging (Figure 2C) [25].

**Figure 4:**
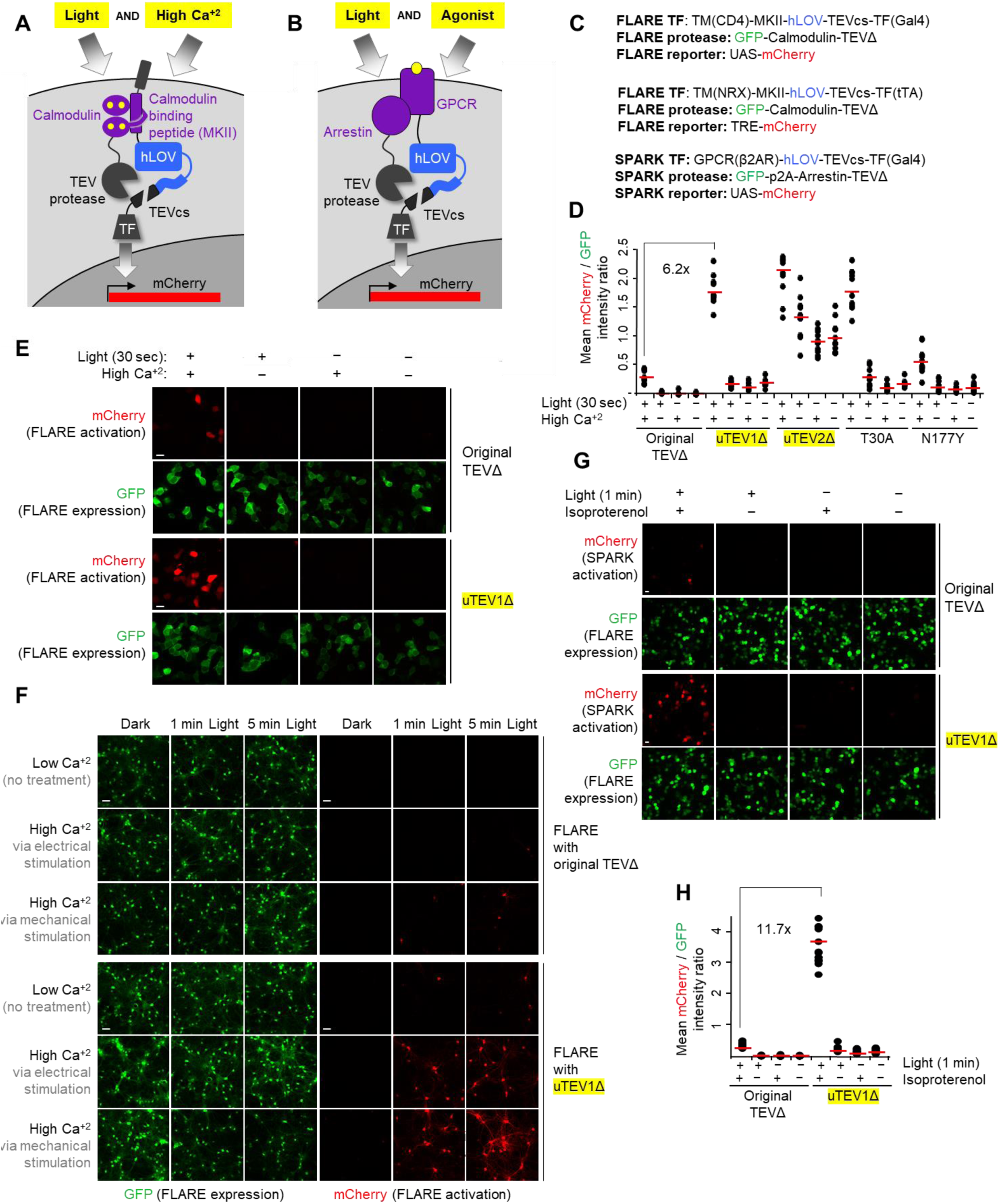
Characterization of evolved low-affinity TEVΔ proteases in mammalian cells and incorporation into FLARE and SPARK tools. A FLARE tool used to integrate cytosolic calcium activity. FLARE is a coincidence detector of blue light and high calcium, with gene expression as the readout [8]. High calcium drives intermolecular complexation between calmodulin and its binding peptide (MKII), which brings TEVΔ protease close to its peptide substrate TEVcs. Blue light is also required to uncage TEVcs. Released TF translocates to the nucleus and drives mCherry expression. B SPARK tool used to integrate GPCR activity. SPARK is a coincidence detector of light and GPCR activity, with gene expression as the readout. Activated GPCR recruits the effector beta-arrestin, which brings TEVΔ protease close to its peptide substrate TEVcs. Blue light is also required to uncage TEVcs. Released TF translocates to the nucleus and drives mCherry expression. C Genetic constructs used for FLARE and SPARK experiments. The first and third set are for HEK293T cells and the second set is for expression in neurons. hLOV is an improved LOV domain described in [11]. p2A is a self-cleaving peptide [36]. D Testing protease mutants using FLARE in HEK293T cells. The indicated protease was incorporated into FLARE as shown in (A) and (C). After transient transfection into HEK293T cells, cells were stimulated with 5 mM CaCl_2_ and ionomycin for 30 sec in the presence or absence of blue light. Eight hours later, mCherry was imaged. Dots indicate quantification of mCherry intensity relative to GFP signal across 10 fields of view per condition (n=10). Red lines indicates the mean of 10 FOVs (**Supplementary Figure 11**). For uTEV1Δ, the light/dark signal ratio is 15, and the high/low Ca^+2^ signal ratio is 12. This experiment was performed two times with similar results. E Sample confocal fluorescence images from the first 8 columns in (D). mCherry reflects FLARE turn-on. GFP reflects expression level of the FLARE tool (protease component). Scale bar, 20 μm. This experiment was performed independently three times with similar results. F uTEV1Δ improves FLARE performance in cultured neurons. Rat cortical neurons were transduced on day 12 with FLARE AAV1/2 viruses. 6 days later (at DIV18), we stimulated the neurons either electrically (3-s trains consisting of 32 1-ms 50 mA pulses at 20 Hz for a total of 1 or 5 min) or mechanically (via replacement of spent media with fresh media of identical composition). The light source was 467 nm, 60 mW/cm2, 10% duty cycle (0.5s light every 5s). 18 hours later, cells were imaged by confocal microscopy (**Supplementary Figure 12**). This experiment was replicated three times. G uTEV1Δ improves SPARK performance in HEK293T cells. SPARK constructs as shown in (C) containing either wild-type TEVΔ or uTEV1Δ were transiently expressed in HEK, and cells were stimulated with 10 μM isoproterenol for 1 min in the presence or absence of blue light. Nine hours later, mCherry was imaged. GFP reflects SPARK expression level. Scale bar, 10 μm. H Quantification of experiment in (G), Dots indicate quantification of mCherry intensity relative to GFP signal across 10 fields of view per condition (n=10). Red lines indicates the mean of 10 FOVs (**Supplementary Figure 13**). For uTEV1Δ, light/dark signal ratio is 22.1, and the +/-agonist signal ratio is 20.7. This experiment was performed two times with similar results.

Wild-type TEVΔ gave an initial turnover rate (*k*_*app*_) of 10×10^−3^ sec^−1^, while the evolved proteases uTEV1Δ and uTEV2Δ gave rates of 54×10^−3^ sec^−1^ and 62×10^−3^ sec^−1^, which are respectively 5.4- and 6.2-fold higher than wild-type TEVΔ. Their differences in actual k_cat_ may be even greater, but we could not measure them by this assay due to our inability to saturate the TEV active sites (to reach *V*_*max*_). Our results, combined with the yeast-based characterization in Figure 2B, suggest that our directed evolution achieved the goal of increasing the catalytic efficiency of TEVΔ while retaining low substrate affinity.

### Characterization of protease sequence-specificity

The mutations in uTEV1Δ and uTEV2Δ border the catalytic triad and are distal to the substrate binding pocket (Figure 1G), making them unlikely to affect the sequence-specificity of TEV. Nevertheless, we wondered whether this extremely valuable and useful property of TEV was affected by our evolved mutations. To evaluate sequence specificity, we turned again to our yeast platform but coupled it to sequencing analysis. We prepared yeast strains expressing TEVcs libraries, caged by LOV and tethered to a transcription factor as shown in Supplementary Figure 5A. The query protease was co-expressed as a fusion to BFP and CRY. After blue light irradiation for 30 minutes, we used FACS to enrich the cells with high Citrine expression. Sequencing of these cells revealed the collection of TEVcs sequences that are capable of being cleaved by the protease of interest. As proof of concept, we mutated specific positions in TEVcs to alanine – P6 (the 6^th^ amino acid N-terminal to the cut site), P3, and P1 – because previous studies have shown that TEV is most sensitive to amino acid changes at these positions [24, 26, 27]. Figure 2F shows that, as expected, mutation of P6, P3, or P1 to alanine largely abolishes TEV recognition.

We then evaluated uTEV1Δ, uTEV2Δ and wild-type TEVΔ sequence specificities using seven TEVcs libraries, each with randomization at one of the positions P6-P1 or P1’ (the residue immediately C-terminal to the cut site) [28]. When we sequenced the libraries post-FACS (Supplementary Figure 5C) we found that the sequence preferences of uTEV1Δ and uTEV2Δ were unchanged from that of wild-type TEVΔ (Figure 2G). [29]

As a further indirect measure of sequence specificity, we evaluated the viability of HEK cells expressing each of the TEV proteases over 3 days. We found that overexpression of the evolved TEVs (both truncated and full-length) did not negatively impact cell health compared to overexpression of wild-type TEV (Supplementary Figure 5D). From these results, we conclude that uTEV1Δ and uTEV2Δ should be useful for cellular applications for which protease orthogonality is a requirement.

### Directed evolution of full-length, high-affinity TEV proteases

While proximity-dependent TEVs are necessary for transcriptional reporters such as FLARE, SPARK, TANGO, and Cal-Light, other applications in biotechnology could benefit from improved *high-affinity* TEVs. We explored the use of our yeast platform for also evolving improved full-length TEV variants. Our selection scheme in Figure 3A differs from the original one (Figure 1A) in that CRY is omitted, and TEVcs is the high-affinity sequence ENLYFQ/S rather than the low-affinity sequence ENLYFQ/M used earlier. This platform was optimized to improve dynamic range (Supplementary Text 2 and Figure 3B), and then applied to an error-prone PCR library of full-length uTEV1 variants. After three rounds of selection, we enriched several mutations that appeared to improve TEV activity (Supplementary Text 3 and Figures 3C-D). The best mutant was uTEV3 (I138T/S153N/T180A), which has 2.8-fold higher k_cat_/K_m_ than wild-type TEV (Figures 3E-F). Interestingly, the improvement comes from a 3-fold decrease in K_m_ rather than an increase in k_cat_ (Figure 3F). Using our yeast-based substrate profiling assay (Supplementary Figure 5), we determined that uTEV3 retains the high sequence-specificity of wild-type TEV (Supplementary Figure 9) and could be a useful reagent for removal of protein affinity tags (Figure 3G). uTEV3 compares favorably to other full-length TEV variants previously engineered by Iverson [17] and Bottomley [30] (Supplementary Text 4).

### uTEV1Δ improves the performance of FLARE and SPARK tools

TEV is utilized in a wide range of biotechnological tools, many of which could benefit from faster protease catalysis. Two such tools are FLARE[8] and SPARK [11, 12], which are caged transcription factors that are activated by the coincidence of blue light and a second stimulus – for FLARE, the second stimulus is elevated cytosolic calcium, while for SPARK, it is a protein-protein interaction (PPI) (Figures 4A and 4B). Both tools convert transient cellular events into stable signals that enable microscopy, manipulation, or genetic selection. How transient an event can be recorded by FLARE or SPARK depends on how many TF molecules can be released per unit time, which, in turn, is limited by protease catalytic rate. Previously, we found that FLARE requires a minimum of 10-15 minutes of light + calcium to give sufficient signal/noise in neuron culture [8]. Similarly, SPARK requires a 10-15 minute “recording” time window to capture a cellular PPI event [11]. We were interested to know whether uTEV1Δ or uTEV2Δ could improve the temporal resolution of FLARE and SPARK tools.

We started by introducing uTEV1Δ, uTEV2Δ, and two other truncated TEV variants into FLARE, and then testing the resulting constructs in HEK 293T cells. Cells were treated with 6 mM CaCl_2_ and 2 μM ionomycin to elevate cytosolic calcium, while blue light was delivered for just 30 seconds. Reporter gene (mCherry) expression was detected by confocal microscopy 8 hours later. Figures 4D-E show that all evolved proteases gave increased mCherry expression compared to wild-type TEVΔ. However, uTEV2Δ was accompanied by high background in the no-light and low-calcium conditions (Figure 4D), perhaps because the eLOV domain is no longer sufficient to fully cage TEVcs against this highly active protease. FLARE with uTEV1Δ gave the best signal-to-background ratio (12.2) and a 6.2-fold improvement over original FLARE with wild-type TEVΔ.

We then moved on to test uTEV1Δ in neuron culture. Here we could elevate cytosolic calcium in a more physiological manner, by using either electrical field stimulation or media replacement, which mechanically stimulates the neurons while providing fresh glutamate. We concurrently delivered blue light at 60 mW/cm^2^ for either 5 minutes or 60 seconds. Figure 4F shows that original FLARE with wild-type TEVΔ gives minimal reporter gene (mCherry) turn-on, consistent with previous observations [11]. By contrast, FLARE incorporating uTEV1Δ shows calcium- and light-dependent reporter gene expression after both 5 minute and 60-second stimulations. Quantitation showed signal-to-noise ratios 27-fold and 16-fold higher, for FLARE containing uTEV1Δ compared to FLARE containing wild-type TEVΔ, at 60 sec and 5 min timepoints, respectively (electrical stimulation conditions). uTEV1Δ therefore improves the performance and temporal resolution of FLARE in neuron culture. (Supplementary Figure 12).

To test uTEV1Δ in SPARK, we selected beta-2-adrenergic receptor (β2AR) and beta-arrestin as our protein-protein interaction pair. Isoproterenol stimulates this interaction, as arrestin is recruited to the GPCR as part of its desensitization pathway. [31] Previously, SPARK required at least 10 minutes of light stimulation to give detectable reporter gene expression in HEK 293T cells. [11] Figure 4G shows that with just 1 minute of isoproterenol + light, mCherry is robustly detected by confocal microscopy. By contrast, original SPARK with uTEVΔ gives a 11.7-fold lower mCherry signal under matched conditions (Figures 4G-H). Hence uTEV1Δ also improves the temporal resolution of the PPI transcriptional tool SPARK.

## Discussion

In this study, we developed a yeast-based platform for the evolution of protease catalytic rate. We used it to improve kinetic parameters for both full-length TEV protease and its truncated, low affinity variant. The latter was then incorporated into the cellular transcriptional reporters FLARE and SPARK to improve the temporal resolution of calcium and protein-protein interaction detection, respectively.

Our directed evolution approach differs in some key respects from previous platforms used to evolve enzyme function. In contrast to selections on the yeast cell surface (used for APEX [32], TurboID [19], split HRP [20], and Iverson’s TEV [17]), our selection takes place in the yeast cytosol, which is more physiologically relevant, and enables us to tie protease catalytic activity to transcription of a fluorescent protein. As a consequence, our signal is amplified, and each selection step is very simple to perform, not requiring any antibody staining. A second feature of our platform is the use of the photosensory LOV domain [33, 8] to cage the TEV cleavage sequence (TEVcs). In doing so, we could modulate the time window available for TEV action on TEVcs, and progressively increase selection stringency. Third, because we used the light-sensitive CRY-CIBN interaction [22] to recruit TEV to its peptide substrate, we could perform selections on low affinity (high *K*_*m*_) proteases, which are required for TANGO, FLARE, Cal-Light and SPARK tools.

The simplicity and modularity of our yeast evolution platform suggest that it could be adapted for other engineering or analysis goals. We showed here that with some small modifications, the system could be used to characterize protease sequence-specificity via sequencing of FACS-enriched clones (Supplementary Figure 5). Alternative strategies for characterizing protease sequence-specificity, using synthetic peptide libraries [34] or N-terminal capture with subtiligase [35], require expensive peptide synthesis or mass spectrometry. Potentially, this platform could also be used to improve the catalytic efficiency of other proteases (as an example, replacement of TEV with TVMV protease in this platform is shown in Supplementary Figure 14), reprogram protease sequence-specificity (an example is shown in Supplementary Figures 15-16), introduce light or analyte-regulation of protease activity, improve/create split proteases, or evolve the properties of photosensory domains.

## ACKNOWLEDGEMENTS

We are grateful to Stanford, the Chan Zuckerberg Biohub, and the Beckman Technology Development Seed Grant for support of this work. FACS was performed at the Koch Institute Flow Cytometry Core (MIT) and at the Stanford Shared FACS Facility. Prof. Wenjing Wang (University of Michigan) provided plasmids and advice. Dr. Lin Ning (Stanford University) provided rat brain tissue. Alex Geoffrey Johnson (Stanford) gave advice on TEV expression, and Nestor Samiylenko helped reproduce some experiments. Dr. Brett Babin, Joshua Yim, and Prof. Matt Bogyo (Stanford University) provided access to their HPLC. Gang Liu (MIT) built the LED box used for blue light irradiation of cells. Maja Djuristic (Stanford University) assisted with electrical stimulation of neurons. M.I.S. was supported by an EMBO long-term post-doctoral fellowship (ALTF 1022-2015).

## AUTHOR CONTRIBUTIONS

M. I. S. performed all the experiments. M. I. S. and A.Y. T. designed the research, analyzed the data, wrote and edited the paper.

The authors declare competing financial interest.

## METHODS

### Cloning

See Plasmid table for a list of genetic constructs used in this study. Each entry lists construct features including promoters, linker sequences, selection markers, epitope tags, etc. For cloning, PCR fragments were amplified using Q5 polymerase (New England BioLabs (NEB)). The vectors were double-digested and ligated to gel-purified PCR products by T4 ligation or Gibson assembly. Ligated plasmid products were introduced by heat shock transformation into competent XL1-Blue bacteria.

### Protease and cleavage site (TEVcs) sequences

All TEV protease constructs used in this study including “wild-type TEV” contain the S219V mutation which inhibits self-proteolysis [23]. All TEVΔ constructs are truncated at position 219V (last C-terminal amino acid is Val). Mutations relative to wild-type TEV in:

uTEV1: S153N

uTEV2: T30A, S153N

uTEV3: I138T, S153N, T180A

Full sequences of each of the above evolved proteases are given in the Plasmid table. In this study we also used two different TEV cleavage site sequences (TEVcs; back slash indicates the proteolysis site):

Low-affinity TEVcs (used for TEVΔ selections and characterization): ENLYFQ/M

High-affinity TEVcs (used for TEV selections and characterization): ENLYFQ/S

### Construction of yeast strains

All strains were derived from Saccharomyces cerevisiae BY4741 (Euroscarf, Johann Wolfgang Goethe-University Frankfurt, Germany). Plasmid transformation or integration in yeast was performed using the Frozen E-Z Yeast Transformation II kit (Zymoprep) according to manufacturer protocols.

S. cerevisiae strains were produced step-wise and propagated at 30 °C in supplemented minimal medium, (SMM; 6.7 g/L Difco nitrogen base without amino acids, 20 g/L dextrose, 0.54 g/L CSM –Ade – His –Leu –Lys –Trp –Ura (Sunrise Science Products)). Transformants were isolated in appropriate selective SD medium by auxotrophic complementation. For yeast strain transformation, we grew cells at 30 °C in YPD containing 10 g/L yeast extract (BD Biosciences, Germany), 20 g/L peptone (BD Biosciences, Germany) and 20 g/L dextrose.

We first obtained the yeast strain containing the reporter gene by integrating (lexA-box)_x_-PminCYC1-Citrine-TCYC1 plasmids (x = 4, 2 or 1) (Addgene plasmids #58434, #58433 and #58432, a gift from Joerg Stelling) into BY4741 [37]. For integration, the plasmid was linearized with PacI. Transformed cells containing the URA3 gene were selected on SMM plates (SMM with 20 g/L agar) and propagated in SMM at 30 °C supplemented with 20 mg/L histidine and 100 mg/L Leucine, producing BY4741-*ura3Δ0*::(lexA-box)4-PminCYC1-Citrine-tCYC1.

With this strain in hand, we proceed to integrate different pRS-derived constructs bearing the membrane-anchored transcription factors. STE2 (Addgene #32171) was a gift from Linda Hicke, promoters/terminators and different TFs were also a gift from Joerg Stelling (Addgene plasmids #58434, #58431, #58438, and #64511) [37]. For integration, plasmids were digested with AscI. Transformed cells containing the LEU2 gene were selected on SMM plates (SMM with 20 g/L agar) and propagated in SMM at 30 °C supplemented with 20 mg/L histidine.

Plasmids containing different TEV protease versions were episomally introduced in a pRSII413 vector which was a gift from Steven Haase (Addgene #35450). Transformed cells containing the HIS3 gene were selected on SMM plates (SMM with 20 g/L agar) and propagated in SMM at 30 °C. For the sequence specificity protease profiling in yeast (Figures 2F-G and Supplementary Figures 5B-C), BY4741-*ura3Δ0*::(lexA-box)4-PminCYC1-Citrine-tCYC1 was also used. In this case, wild type and evolved proteases in full length or truncated forms (fused to mtagBFPII to detect protease expression) were integrated into the Leu2Δ1 locus after digestion with AscI. Transformed cells containing the LEU2 gene were selected on SMM plates (SMM with 20 g/L agar) and propagated in SMM at 30 °C supplemented with 20 mg/L histidine. Next, the cells were transformed with TEVcs plasmid libraries (in pRSII413 backbone with HIS3 as an auxotrophic marker) and selected on SMM plates and propagated in SMM at 30 °C.

See Yeast strain table for a listing of yeast strains used in this study.

### Yeast growth, induction, and light stimulation

Single colonies of transformed cells containing the 3 components (reporter gene, membrane-anchored TF, and protease) were inoculated in 5 mL SSM media and cultured at 30 °C and 220 r.p.m. The fresh saturated culture was diluted 1:20 in fresh media of identical composition and allowed to grow for approximately 6-9 h more until reaching OD600 ∼0.6.

TEV protease expression was induced by inoculating 0.25 mL of a non-saturated yeast culture (OD600 ∼0.6) into 4.75 mL of 10% D/G SMM (SMM medium with 90% of dextrose replaced with galactose) at 30 °C and 220 r.p.m. for 12 h. An aliquot of this culture (around 0.25 mL) was placed in a cuvette at ~4 cm distance from the light source of a MaestroGen UltraBright LED transilluminator (continuous illumination at 470 nm for the indicated time periods). The irradiated sample was transfer to an Eppendorf tube and incubated in a rotator for 6 hours in the dark at 30 °C.

Note: We observed that the activity of the *TDH3* promoter (which controls expression of the membrane-anchored TF) with time and saturation of the yeast culture [38]. Therefore, we were careful to always pay attention to growth times and OD600 values.

### FACS analysis and sorting of yeast populations

Yeast samples, after incubation for 6 h in the conditions described above, were transferred to a 5 mL polystyrene round-bottom tube with 1 mL of DPBS (0.2 mL yeast was diluted into 1 mL of DPBS).

For two-dimensional FACS analysis, we used a LSRII-UV flow cytometer (BD Biosciences) to analyze yeast with 488 nm and 561 nm lasers and 525/50 (for citrine) and 610/20 (for mCherry) emission filters. To analyze and sort single yeast cells, cells were plotted by forward-scatter area (FSC-A) and side-scatter area (SSC-A) and a gate was drawn around cells clustered between 10^4^ – 10^5^ FSC-A and 10^3^ – 10^5^ SSC-A to give population P1. Cells from population P1 were then plotted by side-scatter width (SSC-W) and side-scatter height (SSC-H) and a gate was drawn around cells clustered between 10 – 100 SSC-W and 10^3^ – 10^5^ SSC-H to give population P2. Cells from population P2 were then plotted by forward-scatter width (FSC-W) and forward-scatter height (FSC-H) and a gate was drawn around cells clustered between 10 – 100 FSC-W and 10^3^ – 10^5^ FSC-H to give population P3. Population P3 often represented >90% of the total population analyzed. From population P3, we then plotted mCherry (561 nm laser and 610/20 emission filter) on the x-axis (representing expression level of the protease or TEVcs) and Citrine (488 nm laser and 525/50 emission filter) on the y-axis (representing turn-on of the reporter gene).

To sort yeast populations, we used a BD Aria II cell sorter (BD Biosciences) with the same parameters explained above. From population P3, gates were drawn to collect yeast with the highest activity/expression ratio, i.e., high Citrine/mCherry ratio, but mCherry intensity greater than ~10^2^ to exclude cells not expressing the protease or TEVcs. BD FACSDIVA software was used to analyze all data from FACS sorting and analysis.

### Directed evolution of truncated, low-affinity TEVΔ in yeast

For the directed evolution of TEVΔ, which comprises amino acids 1-219 of the wild type TEV protease, three libraries were generated using TEVΔ-S219V as the starting template. To perform error-prone PCR on this template, we combined 100 ng of the template plasmid (GalP-mCherry-CRY2PHR-TEVΔ in pRSII413) with 0.4 μM forward and reverse primers that anneal to the sequences just outside the 5′ and 3′ ends of the gene encoding TEVΔ, 2 mM MgCl_2_, 10 units of Taq polymerase (NEB), 0.2 mM of regular dNTPs, 1× Taq polymerase buffer (NEB) and 2 μM or 20 μM each of the mutagenic nucleotide analogs 8-oxo-2′-deoxyguanosine-5′-triphosphate (8-oxo-dGTP) and 2′-deoxy-p-nucleoside-5′-triphosphate (dPTP) in a total volume of 100 μL. We used the following conditions to produce varying levels of mutagenesis:

Library 1: 2 μM 8-oxo-dGTP, 2 μM dPTP, 10 PCR cycles

Library 2: 2 μM 8-oxo-dGTP, 2 μM dPTP, 20 PCR cycles

Library 3: 20 μM 8-oxo-dGTP, 20 μM dPTP, 10 PCR cycles

Forward primer: 5′-ggtggaagtggatcaggcagcggtggatctggcagcggaaagcttggttccggg-3’

Reverse primer: 5′-ggagggcgtgaatgtaagcgtgacataactaattacatgactcgagctatta-3’

The PCR was performed with an annealing temperature of 58 °C per cycle. The PCR products were gel-purified, then re-amplified under regular conditions for another 30 cycles with 0.4 μM forward and reverse primers that introduce ∼45 bp of overlap with both ends of the vector.

Separately, we prepared digested vector. We linearized the pIIRS-413:GalP-mCherry-CRY2PHR-TEVΔ-tCYC1 plasmid by digesting with HindIII-HF and XhoI restriction enzymes overnight at 37 °C. These enzymes digest the gene just upstream and downstream of the TEVΔ gene. The linearized vector backbone was purified by gel extraction. We then combined 1 μg of linearized vector with 4 μg of mutagenized TEVΔ PCR product from above, and concentrated using pellet paint (Millipore) according to the manufacturer’s protocols. The DNA was precipitated with ethanol and sodium acetate, and resuspended in 10 μL ddH2O.

Fresh electrocompetent BY4741 yeast containing the reporter gene (lexA-box)_4_-PminCYC1-Citrine-tCYC1 integrated in the *ura3Δ0* locus, and the optimized TF TDH3:STE2*Δ-*CIBN-BFP-eLOV-TEVcs(ENLYFQ/M)-LexAVP16-tCYC1 in the Leu2Δ1 locus, were prepared.

Fresh electrocompetent BY4741 yeast containing the reporter gene (lexA-box)_4_-PminCYC1-Citrine-tCYC1 integrated in the *ura3Δ0* locus, and the optimized TF TDH3:STE2*Δ-*CIBN-BFP-eLOV-TEVcs(ENLYFQ/M)-LexAVP16-tCYC1 in the Leu2Δ1 locus, were prepared. Yeast were passaged at least two times before this procedure to ensure that the cells were healthy. We used 2-3 mL of an overnight culture to inoculate 100 mL of YPD media. The culture was grown with shaking at 220 r.p.m. at 30 °C for 6–8 h until the OD600 reached 1.5–1.8. Yeast were then harvested by centrifugation for 3 min at 3,000 r.p.m. and resuspended in 50 mL of sterile 100 mM lithium acetate in water by vigorous shaking. Fresh sterile DTT (1 M stock solution, made on the same day) was added to the yeast cells to a final concentration of 10 mM. The cells were incubated with shaking at 220 r.p.m. for 12 min at 30 °C (necessary to ensure adequate oxygenation). Then yeast were pelleted at 4 °C by centrifugation at 3,000 r.p.m. for 3 min and washed once with 25 mL ice-cold sterile water, pelleted again, and resuspended in 1 mL ice-cold sterile water.

The concentrated mixed DNA from above was combined with 250 μL of electrocompetent yeast placed into a Gene Pulser Cuvette (BIO-RAD, catalog 3165-2086) prechilled in ice and then electroporated using a Bio-Rad Gene pulser XCell with the following settings: 500-V, 15-ms pulse duration, one pulse only, 2-mm cuvette. The electroporated cells were immediately rescued with 2 mL pre-warmed YPD media and then incubated at 30 °C for 2 h without shaking. Cells were vortexed briefly, and 1.99 mL of the rescued cell suspension was transferred to 100 mL of SMM medium supplemented with 50 units/mL penicillin and 50 μg/mL streptomycin and grown for 2 days at 30 °C. The remaining 10 μL of the rescued cell suspension was diluted 100×, 1000×, 10000×, and 100000×; 20 μL of each dilution was plated on SMM plates and incubated at 30 °C for 3 days. After 3 days, each colony observed in the 100×, 1000×, 10000×, or 100000× segments of plates will correspond to 10^4^, 10^5^, 10^6^, or 10^7^ transformants in the library, respectively. The culture was grown at 30 °C with shaking at 220 r.p.m. for 1 d, before induction of protein expression and positive selection as described below (“Yeast selections”).

The transformation efficiency of our TEVΔ library into BY4741-*ura3Δ0*::(lexA-box)_4_-PminCYC1-Citrine-tCYC1/Leu2Δ1::TDH3:STE2*Δ-*CIBN-BFP-eLOV-TEVcs(ENLYFQ/M)-LexAVP16-tCYC1 pRSII413-HIS3 was determined to be ~4 × 10^7^. DNA sequencing of 24 individual clones showed that each clone had 0–8 amino acids changed relative to the original TEVΔ template.

For the directed evolution of full-length TEV, we prepared an error-prone PCR library based on the template uTEV3 in the same manner described above. The following primers were used for amplification: Forward primer: 5′-gtggaggcggtagcggaggcggagggtcggctagcggcagcggaaagcttggttccggg-3’ Reverse primer: 5′-ggagggcgtgaatgtaagcgtgacataactaattacatgactcgagctatta-3’ The transformation efficiency of the uTEV3 library into BY4741-*ura3Δ0*::(lexA-box)_2_-PminCYC1-Citrine-tCYC1/Leu2Δ1::TDH3:STE2*Δ-*CIBN-BFP-eLOV-TEVcs(ENLYFQ/S)-LexAVP16-tCYC1, pRSII413-HIS3 was determined to be ~31 × 10^7^. DNA sequencing of 24 individual clones showed that each clone had 0– 7 amino acids changed relative to the original template.

### Yeast selections

For each round of selection, we input ~10-fold more yeast cells than the estimated library size. For the first round, library size was estimated by the transformation efficiency of the initial protease library. For subsequent rounds, library size was taken to be the number of yeast cells collected during the previous sort.

Yeast cells transformed with the TEVΔ library in pRSII413 (prepared as described in “Generation of TEV mutant libraries and transformation into yeast’’) were induced by transferring them to 1:10 SMM-D/G media, and growing the cells for 12 h at 30 °C with shaking at 220 r.p.m. For the first round of selection, 10 mL of yeast culture (at OD_600_ ∼1.5; note that OD_600_ ∼1 corresponds to roughly 3 × 10^7^ yeast cells/mL) were placed in cuvettes at ~4 cm distance from the light source of a MaestroGen UltraBright LED transilluminator (continuous illumination) at 470 nm for 8 min.

After irradiation, samples were split between two culture tubes, each containing 1 mL D/G SMM, and incubated for 6 h at 30 °C with shaking at 220 r.p.m. Cultures were spun down at 3,000 r.p.m. for 3 min and resuspended in a total of 5 mL PBS-B (sterile phosphate-buffered saline supplemented with 0.1% BSA).

FACS sorting was carried out as described above (“yeast selections”). Cells collected from the FACS instrument were immediately placed in a 30 °C incubator with shaking at 220 r.p.m in SSM + 1% pen-strep. Cells were grown for 1-2 days until saturation. Yeast cells were then passaged in this manner at least two more times prior to the next round of selection.

For the TEVΔ evolution, cells were collected from each round as follows:

Round 1: 1.5 % of cells collected (6 × 10^6^ cells)

Round 2: 1 % of cells collected (6 × 10^5^ cells)

Round 3: 0.5% of cells collected (3 × 10^4^ cells)

After the third round of selection, yeast were collected as before, and 1 mL of the growing culture (at OD600 ~1.2) was removed for DNA extraction using the Zymoprep yeast Plasmid Miniprep II kit according to manufacturer protocols. After plasmid transformation into XL1B bacteria, single colonies were grown overnight at 37 °C with shaking at 220 r.p.m. Bacterial cultures were spun down at 6000 rpm for 6 min and plasmid was extracted using the QIAprep Spin Miniprep Kit according to manufacturer protocols. TEV mutants were analyzed by Sanger sequencing. The sequencing primer used was: 5’-cgcagattatgatcggagcagcgccg-3’.

### Profiling protease sequence specificity in yeast (related to Figures 2G and Supplementary Figures 5B-C)

To prepare the TEVcs libraries for profiling protease substrate specificity, we digested the plasmid RSII413:GalP-STE2Δ-mCherry-CIBN-PIF6-eLOV-TEVcs(ENLYFQM)-LexAVP16-tCYC1 overnight with BamHI-HF and HindIII-HF enzymes at 37 °C. The linearized vector was purified by gel extraction. Separately, we carried out PCR on the template plasmid RSII413:GalP-STE2Δ-mCherry-CIBN-PIF6-eLOV-TEVcs(ENLYFQM)-LexAVP16-tCYC1 using the forward primer 5’-gttggaaagcaataaacatgttgacgggggatcc-3’ and one of the following reverse primers:

R-P6: 5’-gcctggccgttaacgctttcataagcttcccgcccatctggaagtagagatt**NNN**cttagcggcttc −3’

R-P5: 5’-gcctggccgttaacgctttcataagcttcccgcccatctggaagtagag**NNN**ttccttagcg −3’

R-P4: 5’-gcctggccgttaacgctttcataagcttcccgcccatctggaagta**NNN**attttccttagc −3’

R-P3: 5’-gcctggccgttaacgctttcataagcttcccgcccatctggaa**NNN**gagattttccttag −3’

R-P2: 5’-gcctggccgttaacgctttcataagcttcccgcccatctg**NNN**gtagagattttcc −3’

R-P1: 5’-gcctggccgttaacgctttcataagcttcccgcccat**NNN**gaagtagagattttc −3’

R-P1’: 5’-gcctggccgttaacgctttcataagcttcccgcc**NNN**ctggaagtagag −3’

PCR products were gel-purified using QIAprep Spin Miniprep Columns, then combined with purified vector and Gibson assembly master mix. The product was transformed into competent XL1-Blue bacteria. After 20 h, colonies were harvested with 5 mL of Luria Broth (LB) supplemented with 100 μg/mL ampicillin and grown overnight at 37 °C with shaking at 220 r.p.m. Bacterial cultures were spun down at 6000 rpm for 6 min and plasmid was extracted with QIAprep Spin Miniprep Kit according to manufacturer protocols. The TEVcs libraries were transformed into yeast containing integrated reporter gene and TEV proteases, using protocols described above under “Construction of yeast strains”. After 48 h, single colonies appeared in SMM-plates and were harvested with 5 mL of SMM and grown overnight at 30 °C with shaking at 220 r.p.m.

TEVcs library expression was induced in these cells by transferring them to 1:10 SMM-D/G media, and growing the cells for 12 h at 30 °C with shaking at 220 r.p.m. 1 mL of yeast culture was placed in a cuvette at ~4 cm distance from the light source of a MaestroGen UltraBright LED transilluminator (continuous illumination at 470 nm for the indicated time periods).

After irradiation, samples were transferred into two culture tubes and incubated for 6 h at 30 °C with shaking at 220 r.p.m. Cultures were spun down at 3,000 r.p.m. for 3 min and resuspended with 1 mL of PBS-B (sterile phosphate-buffered saline supplemented with 0.1% BSA). FACS sorting was performed to select cells with high Citrine/mCherry ratio, as described above under “FACS analysis and sorting of yeast populations”.

Cells collected from the FACS instrument were immediately placed in a 30 °C incubator with shaking at 220 r.p.m in SSM + 1% pen-strep. Cells were grown for 1-2 days until saturation. Then 1 mL of the culture was removed for DNA extraction using the Zymoprep yeast Plasmid Miniprep II (Zymo Research) kit. After plasmid transformation into XL1B bacteria, single colonies were grown overnight at 37 °C with shaking at 220 r.p.m. Bacterial cultures were spun down at 6000 rpm for 6 min and plasmid was extracted with QIAprep Spin Miniprep Kit. Clones were analyzed by Sanger sequencing using the primer 5’-ggtgccatcacaaatctcggggacacgc-3’.

### Fluorescence microscopy of cultured cells

Confocal imaging was performed on a Zeiss AxioObserver inverted confocal microscope with 10× air and 40× oil-immersion objectives, outfitted with a Yokogawa spinning disk confocal head, a Quad-band notch dichroic mirror (405/488/568/647), and 405 (diode), 491 (DPSS), 561 (DPSS) and 640-nm (diode) lasers (all 50 mW). The following combinations of laser excitation and emission filters were used for the fluorophores: eGFP/citrine (491 laser excitation; 528/38 emission), mCherry (561 laser excitation; 617/73 emission), and differential interference contrast. All images were collected and processed using SlideBook (Intelligent Imaging Innovations).

### HEK 293T cell culture and transfection

HEK 293T cells from ATCC with fewer than 20 passages were cultured as monolayers in media composed of a 1:1 mixture of DMEM (Dulbecco’s Modified Eagle medium, Gibco) and MEM (Minimum Essential Medium Eagle) supplemented with 10% (v/v) FBS (Fetal Bovine Serum, Sigma) and + 1% (v/v) pen-strep at 37 °C under 5% CO2. For imaging at 10× magnification, we grew the cells in plastic 48-well plates that were pretreated with 50 μg/mL human fibronectin (Millipore) for at least 10 min at 37 °C before cell plating (to improve cell adherence). For imaging at 40× magnification, we grew cells on 7 × 7 mm glass cover slips placed inside 48-well plates. The coverslips were also pretreated with 50 μg/mL human fibronectin for at least 10 min at 37 °C before cell plating. Cells were transfected at 60–90% confluence with 1 mg/mL PEI max solution (polyethylenimine HCl Max pH 7.3).

For FLARE and SPARK experiments in HEK, cells were transfected in 48-well plates, and each well received a DNA mixture consisting of: 20 ng UAS-mCherry plasmid; 20 ng protease plasmid; and 50– 100 ng of membrane-anchored TF plasmid, along with 0.8 μL PEI max in 10 μL serum-free MEM media for 15 min at room temperature. DMEM/MEM with 10% FBS (100 μL) was then mixed with the DNA-PEI max solution and incubated with the HEK cells for 15-18 h before further processing.

FLARE plasmids used: P55, P56, and one of: P57-P61 (from Plasmid table) SPARK plasmids used: P55, P69, and one of: P70/P71 (from Plasmid table)

### FLARE and SPARK experiments in HEK 293T

HEK 293T cells expressing FLARE constructs were processed 15 h post-transfection. To elevate cytosolic calcium, 100 μL of ionomycin and CaCl_2_ in complete growth media was added gently to the top of the media within a 48-well plate to final concentrations of 2 μM and 6 mM, respectively. For low Ca^2+^ conditions, 200 μL of complete growth media (with no added Ca^+2^) was added. After incubation for the indicated times, the solution in each well was removed and the cells were washed once and then incubated with 200 μL complete growth media.

When Ca^2+^ stimulation was coincident with light illumination, one 48-well plate of HEK 293T cells was placed on top of a custom-built LED light box that delivers 467-nm blue light at 60 mW/cm^2^ intensity and 33% duty cycle (2 s of light every 6 s). Cells were irradiated on the blue LED light box for the indicated time periods. For the dark condition, HEK 293T cells were wrapped in aluminum foil. Afterwards, media in each well was removed, and incubated with 250 μL complete growth media. HEK 293T cells were then incubated in the dark at 37 °C for 8-9 h and imaged right away.

For SPARK experiments, HEK 293T cells were also processed 15 h post-transfection. Complete growth media containing isoproterenol was added gently to the top of each well in a 48-well plate of transfected HEK cells, to a final concentration of 10 μM. For the no-agonist control, 200 μL of complete growth media (lacking agonist) was added. After stimulation for the indicated time periods, the solution in each well was removed, and then 200 μL complete growth media was added. Blue light stimulation was performed as described above for FLARE. Cells were analyzed by microscopy or platereader 8-9 hours post-stimulation.

For Figures 4F and Supplementary Figures 11, 12 and 13, FLARE and SPARK samples were imaged at 10x magnification directly in 48-well plates. More than ten fields of view were typically acquired for each condition. To quantify FLARE or SPARK turn-on, we created a mask based on GFP (which reflects FLARE/SPARK protease expression). Within this mask, we calculated the mean mCherry intensity (background-corrected). These values are reported in the graphs.

### Production of AAV virus supernatant for neuron transduction

AAV virus supernatant was used to transduce neuron cultures for FLARE experiments. To generate viruses, HEK 293T cells were transfected in a T25 flask, at 70–90% confluence. For each virus, we combined 0.875 μg of viral DNA (plasmid P62-P65, from the Plasmid Table), 0.725 μg AAV1 serotype plasmid (plasmid P72), 0.725 μg AAV2 serotype plasmid (plasmid P73), and 1.75 μg helper plasmid pDF6 (plasmid P74) with 20 μL PEI max and 200 μL serum-free DMEM [39]. The transfection mix was incubated for 15 min at room temperature, and added to 5 mL of complete media. The media of the T25 flask was replaced with the mixture. After incubation for 48 h at 37 °C, the supernatant (containing secreted AAV virus) was collected and filtered through a 0.45-μm syringe filter (VWR). AAV virus was aliquoted into sterile Eppendorf tubes (0.5 mL/tube), flash frozen in liquid nitrogen and stored at −80 °C.

### Rat cortical neuron culture

Cortical neurons were harvested from rat embryos euthanized (we have complied with all relevant ethical regulations), at embryonic day 18 and plated in 24-well plates as previously described [40], but without glass cover slips. At DIV4, 300 μL media was removed from each well and replaced with 500 μL complete neurobasal media (neurobasal (Gibco) supplemented with 2% (v/v) B27 supplement (Life Technologies), 1% (v/v) Glutamax (Life Technologies), 1% (v/v) penicillin-streptomycin (VWR, 5 units/mL of penicillin and 5μg/mL streptomycin), and 10 μM (C_f_) 5-fluorodexoyuridine (FUDR, Sigma-Aldrich) to inhibit glial cell growth. Subsequently, approximately 30% of the media in each well was replaced with fresh complete neurobasal media every 3 days. Neurons were maintained at 37 °C under 5% CO2.

### Viral transduction of cortical neuron cultures, stimulation, and data analysis

A mixture of AAV viruses, prepared as above, encoding FLARE components as shown in Figure 4C, was added to cultured neurons between DIV11-12. Typical viral supernatant quantities used were 100 μL of each viral component, added to each well of a 24-well plate, where each well already contained 1.5 mL of complete neurobasal media. After incubation for 3 days at 37 °C, 30% of the media in each well was replaced with fresh complete neurobasal media.

After viral transduction, neurons were grown in the dark, wrapped in aluminum foil, and all subsequent manipulations were performed in a dark room with red light illumination to prevent unwanted activation of the LOV domain. Six days post-transduction (at DIV 17-18), neurons were stimulated in the presence or absence of blue light. To elevate cytosolic Ca^2+^ we used either electrical stimulation or media replacement. For the latter, 30% of the media in each well was replaced with fresh complete neurobasal media of identical composition. After this treatment for 60 sec or 5 min, the saved old culture media was returned to the wells. We found that this improved the health of the neurons. For the low-calcium condition, neurons were not treated (no media change).

For electrical stimulation, we used a Master 8 device (AMPI) to induce trains of electric stimuli. A stimulator isolator unit (Warner Instrument, SIU-102b) was used to provide constant current output ranging from 10–100 mA. Platinum iridium alloy (70:30) wire from Alfa-Aesar was folded into a pair of rectangles (0.7 cm × 1.5 cm) and placed right above the neurons on the edge of the well to act as electrodes. We used 3-second trains, each consists of 32 1-ms 48 mA pulses at 20 Hz, lasting for a total of 60 sec or 5 min. Prior to performing FLARE experiments, we checked each method of stimulation and the quality of our neuron cultures by performing GCaMP5f real-time calcium imaging (data not shown).

For blue-light irradiation, neuron plates were placed on top of the custom-built LED light box described above (“FLARE and SPARK experiments in HEK 293T”) and irradiated with 467-nm blue light at 60 mW/cm^2^ and 10% duty cycle, 0.5 s of light every 5 s. For the dark condition, neurons were wrapped in aluminum foil. Imaging was performed 18 hours later.

For imaging, ten fields of view were collected per condition. For each field of view, a mask was created to encompass regions with GFP expression (reflecting FLARE protease expression). In these masked regions, the mean mCherry fluorescence intensity was calculated, and background was subtracted. These mean mCherry intensity values were calculated individually for 10 fields of view per condition and plotted in a bar plot.

### Bacterial expression and purification of TEV proteases

For bacterial expression and purification of truncated TEVΔ proteases, we cloned the TEV genes into the pRK793 vector, derived from Addgene plasmid #8827 (bacterial expression plasmid for wild-type TEV, from David Waugh).

Following the published protocol from David Waugh [41], competent BL21-CodonPlus(DE3)-RIPL *E. coli* were transformed with the TEV expression plasmids by heat shock. Cells were then grown in TB media (1 L) containing 100 mg/L ampicillin at 37 °C and 220 r.p.m. until OD600 ~0.6. Protein expression was induced with 1 mM (C_f_) IPTG, then cultures were shifted from 37 °C to 25 °C during the induction period to maximize the yield of soluble TEV protease. After overnight growth at 220 r.p.m. and room temperature, the bacteria were pelleted by centrifugation at 6,000 r.p.m. for 6 min at room temperature, the supernatant was discarded, and the pellet was stored at −80 °C.

The frozen pellet was thawed on ice in 50 mL of lysis buffer (50 mM sodium phosphate (pH 8.0), 200 mM NaCl, 10% Glycerol, and 25 mM imidazole) with 1 tablet of complete protease inhibitory (Roche). The pellet was solubilized by pipetting up and down, then the lysate was transferred to a small metal beaker pre-chilled on ice and sonicated using a Misonix sonicator (20 s on, 60 s off, for a total of 6 min on). The sonicated lysate was clarified by centrifugation for 15 min at 11,000 rpm and the supernatant transferred into a 50 mL conical, where it was incubated with 2 mL] Ni-NTA agarose bead slurry (QIAGEN) for 10 min at 4 °C. The slurry was placed in a gravity column and washed with 50 mL of lysis buffer. The protein was eluted with elution buffer (50 mM sodium phosphate (pH 8.0), 200 mM NaCl, 10% Glycerol, and 250 mM imidazole). The purity was analyzed by SDS-PAGE and Coomassie Blue staining.

The eluted samples were dialyzed overnight in 40 mM Tris-HCl (pH 7.5), 200 mM NaCl, 2 mM EDTA, 0.2% Triton X-100, 4 mM beta-mercaptoethanol at 4 °C using a Slide-A-Lyzer Dialysis Cassette (Extra Strength) 10,000 MWCO (Thermo). The dialyzed sample was concentrated using Amicon® Ultra - 15 Centrifugal Filter Units −10,000 NMWL. After concentration, 20% v/v of glycerol was added, and the samples were flash frozen and stored at −80 °C.

For expression and purification of full-length TEV proteases, we cloned genes into a different expression vector - pYFJ16 – which contains an MBP (maltose binding protein) tag in addition to a His6 tag at the N-terminus. The protocol for expression and purification was the same as described above for truncated TEVΔ proteases.

### Cloning, expression, and purification of MBP-TEVcs-GFP

The protease substrate MBP-TEVcs-eGFP (where TEVcs is either ENLYFQ/S or ENLYFQ/M) in pYFJ16 vector was expressed in BL21-CodonPlus(DE3)-RIPL *E. coli*. Cells were grown, induced, and lysed as described above for the TEV proteases. Ni-NTA purification was carried out in the same way. Bright yellow protein fractions were collected and transferred to a centrifugal filter Amicon Ultra-15 and exchanged 3 times into ice-cold 50 mM Tris-HCl buffer (pH 8.0) containing 10% glycerol, 1 mM EDTA, and 2 mM of DTT.

### TEV kinetic assays

The substrate protein MBP-TEVcs-GFP was combined at different concentrations with 100 nM of the indicated protease in 50 mM Tris-HCl buffer (pH 8.0) containing 10% glycerol, 1 mM EDTA, and 2 mM DTT (freshly prepared) at 30 °C. Proteolysis was terminated at various timepoints by mixing the reaction mixtures with SDS-PAGE protein loading buffer and immediately flash freezing in liquid nitrogen. The reactions were then analyzed by SDS–PAGE at 4 °C. Gel fluorescence images were acquired on a Typhoon 9410 instrument, and band intensities were quantified by ImageJ software relative to reference standards of known concentration. We calculated initial reaction velocities at substrate consumption <25%. Data was fit to a Michaelis-Menten enzyme kinetics model with center values representing the mean and error bars representing the standard deviation of three technical replicates.

In Figure 3G, the MBP-TEVcs-GFP concentration was 0.72 mg/ml (10 uM), and the protease concentration was 60 nM. At higher concentrations of protease or substrate, the difference between wild-type TEV and uTEV3 performance is not as obvious.

### Data availability

Additional data beyond that provided in the Figures and Supporting Information are available from the corresponding author upon request.

