## Supporting Information for "Directed evolution improves the catalytic efficiency of TEV protease"

**Supplementary Text 1: Optimization of TEVΔ yeast selection platform**

Using C-terminally truncated, low-affinity wild-type TEV (TEVΔ219 [1], or "TEVΔ") as our starting template, we optimized a number of features of the platform. We found that truncation of the STE2-based plasma membrane anchor (Supplementary Figure 1A) [2] and incorporation of a BFP linker both increased Citrine signal, likely because of improved membrane trafficking of the TF construct. We also increased expression of the membrane-anchored TF by optimizing its promoter [3] (Figure 1C), incorporated a terminator [4] (Figure 1C), and tested alternative activation domains (Supplementary Figure 1C). By performing a time course, we determined that 6 hours was sufficient to reach maximal Citrine expression post light-induction of TF release (Supplementary Figure 1D). Omission of either light or CRY abolished the Citrine signal (Figure 1D).

**Supplementary Text 2: Optimization of full-length TEV yeast selection platform**

While establishing the yeast platform for evolution of full-length, high-affinity TEV proteases, we initially observed signal saturation due to the high affinity between wild-type full-length TEV and its TEVcs. To reduce the sensitivity, and thereby increase the dynamic range of our platform, we experimented with alternative TFs and promoters (Supplementary Figure 6B). We found that by reducing the number of LexA boxes in the promoter of our reporter gene, we could diminish the Citrine response to wild-type TEV [5]. We then tested the mutations enriched in our previous selection (Figure 3B and Supplementary Figure 6C) in the context of full-length TEV. We found that the S153N (from uTEV1Δ) improved activity and therefore incorporated this mutation in our starting template (full-length uTEV1).

**Supplementary Text 3: Directed evolution of full-length TEV**

Using the selection scheme in Figure 3A, we induced protease expression with galactose 12 hours prior to blue light irradiation for 45 seconds (round 1), 30 seconds (round 2), or 5 seconds (round 3). The yeast populations after each round were compared in Figure 3C. The selection clearly enriched activity, but there was little difference between light and dark conditions (Figure 3C and Supplementary Figure 7A), suggesting that eLOV does not effectively cage the high-affinity TEVcs from full-length protease under these conditions. Even without the advantage of light-based kinetic selection as in Figure 1A, however, we nevertheless enriched mutations that improved the catalytic efficiency of full-length TEV. Sequencing of round 3 showed that the mutations S135F, I138T and T180A had been enriched (Supplementary Figure 7B). Assays on individual mutants in yeast showed that all three mutations contributed to improved TEV activity (Supplementary Figure 8A), and combining the mutations had an additive effect (Supplementary Figure 8B). The best final clone, uTEV3, combines these three beneficial mutations (I138T/S153N/T180A).

**Supplementary Text 4: Comparison to previous evolved TEVs**

Previously, Iverson et al. used directed evolution in the yeast secretory pathway to evolve a full-length TEV mutant (G79E/T173A/S219V) with higher activity than wild-type TEV [6]. In another study, a TEV mutant (L56V/S153G/S219V) was designed *in-silico* and shown to exhibit improved solubility and stability [7, 8]. We compared these TEV variants to our own, using our transcriptional assay in the yeast cytosol (Figure 3A). Supplementary Figure 10A shows that uTEV3 is more active than both the previous full-length TEVs. We also performed a comparison in the context of truncated TEVΔ for proximity-dependent cleavage. Supplementary Figure 10B shows that uTEV2Δ was the most active (against the low-affinity TEVcs ENLYFQ/M), while the Iverson mutant (truncated form) and wild-type TEVΔ exhibited comparable activity. In the context of FLARE, the Iverson mutant (truncated form) also performed more poorly than uTEV1Δ (Supplementary Figure 11C). Note that the Iverson mutant was not evolved against the low-affinity TEVcs and likely has lower $k_{cat}$ towards it than do uTEV1Δ and uTEV2Δ.

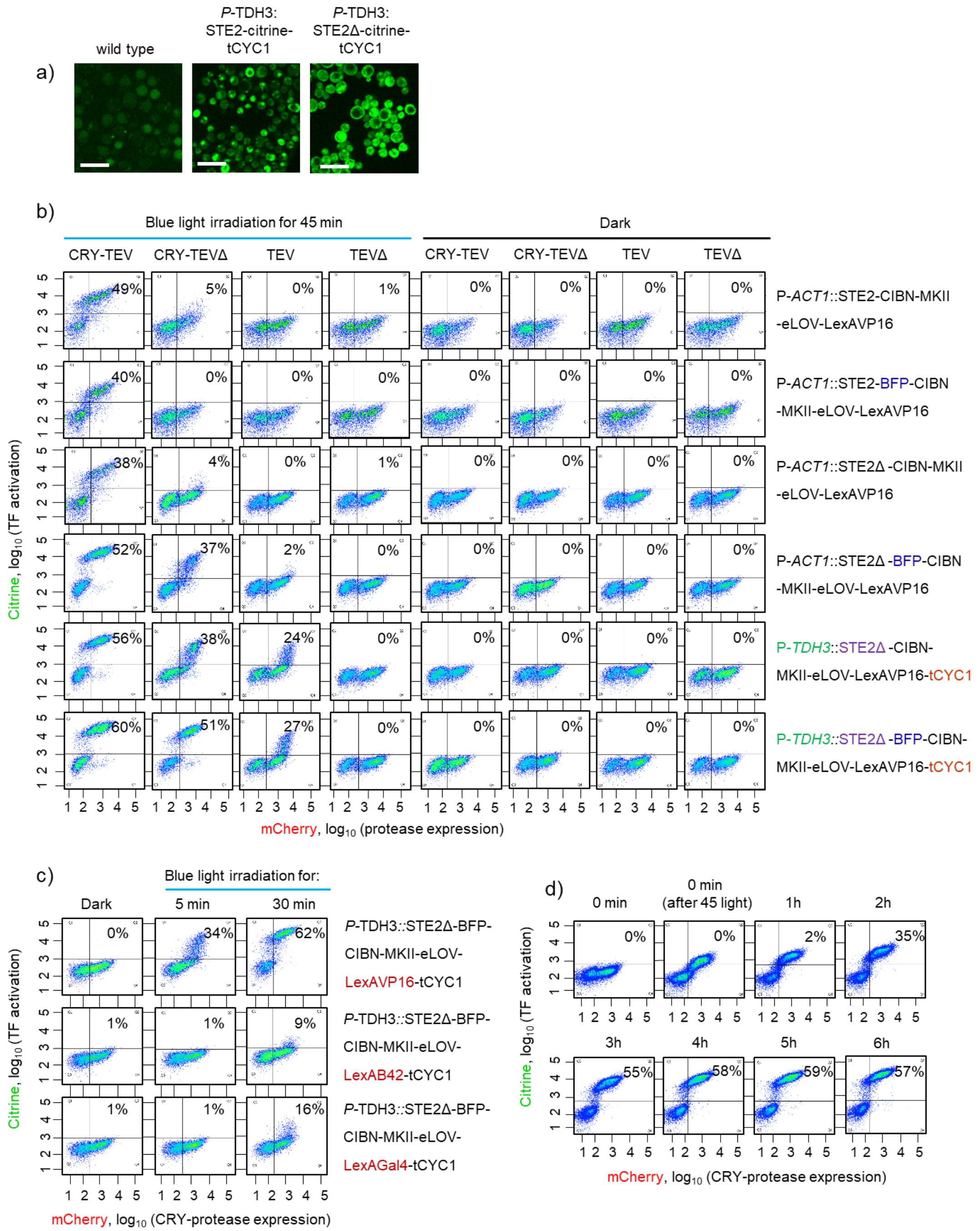


**Supplementary Figure 1. Optimization of membrane-anchored transcription factor for yeast directed evolution.** Related to Figure 1C. (**A**) BY4741 yeast constitutively expressing STE2-citrine or STE2Δ(1-300)-citrine; the latter have much improved surface localization. Scale bars, 10 μm (**B**) Left four columns: FACS plots showing yeast cells 6 hours after 45-minute blue light irradiation. Percentages reflect fraction of cells with Citrine signal (cells that release TF to drive Citrine expression) and are given in the table in Figure 1C. Right four columns: Control cells without light exposure. Each condition performed twice, n = 10,000 cells. (**C**) Optimization of the LexA transcriptional activator. Yeast co-expressing the TF and mCherry-CRY-TEVΔ were analyzed by FACS 6 hours after variable amounts of blue light exposure. Percentages reflect the fraction of Citrine-positive cells. (**D**). Optimizing the time for reporter transcription and translation. FACS plots collected at various timepoints after 45-min blue light exposure to induce TF release. Percentages give fraction of cells in Q1 + Q2 quadrants. Each plot representative of two replicates, n = 20,000 cells. We selected 6 hours as our expression time window.

a)

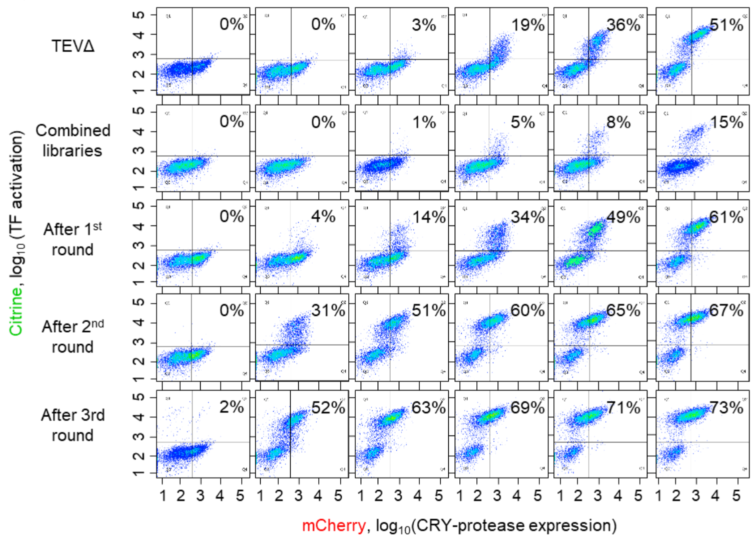


b)

| | | | | | | | | | | | | | | | | | | | |
|---|---|---|---|---|---|---|---|---|---|---|---|---|---|---|---|---|---|---|---|
| #1 | | | | | | S31W | | | | | | | | | | | | | |
| #2 | | | | | T30A | | | | | | | | | | | | | S153N | |
| #3 | | | | | | | | | L56S | | | | | | | | | | N177Y |
| #4 | | | K67E | K89E | | | | | | | | | | | | | | | |
| #5 | | | | | | | | | | | | | | | | | | | N177Y |
| #6 | | | | | | | | | | | | | | | | | | | N177Y |
| #7 | | | | | | | | T54A | | | | | | | | | | | N177Y |
| #8 | | | | | | S31W | | | | | | | | | | | | | |
| #9 | | | | | | | | | | F64L | | | | | | | | | N177Y |
| #10 | | | | | | S31W | | | | | | | | | | | | | |
| #11 | | S16I | | | | | R50G | | | | D90G | | | | | | | | N177Y |
| #12 | | | | | | | | | | | | | | | | | | S153N | |
| #13 | | | | | T30A | | | | | | | | | | | | | S153N | |
| #14 | | | | | | | | | | | | | | | | | | | N177Y |
| #15 | | | | | T30I | | | | | | | | | | | D136V | | | |
| #16 | | | | | | | | | | | | | | | | | | | N177Y |
| #17 | | | | | | | | | | | | I100V | | | | | | | |
| #18 | | | | | | | | | | | | | | | | | | | |
| #19 | | | | | | | | | | | | | | | | | | S153N | |
| #20 | | | | | | | | | | | | | | | S135F | | | | N177Y |
| #21 | | | | | | | | | | | | | | | | | I138T | | |
| #22 | | | | | | S31W | | | | | | | C101R | T114A | | | | | |
| #23 | S3F | | | | | | | | | | | | | | | | | | N177Y |
| #24 | | | | | T30A | | | | | | | | | | | | | | |

**Supplementary Figure 2. Analysis of selected yeast populations and TEV clones.** Related to Figure 1F. (**A**) Same as Figure 1F, but more conditions are shown. (**B**) Sequencing after round 3. 24 clones were sequenced and mutations found in each, relative to original TEVΔ, are shown.

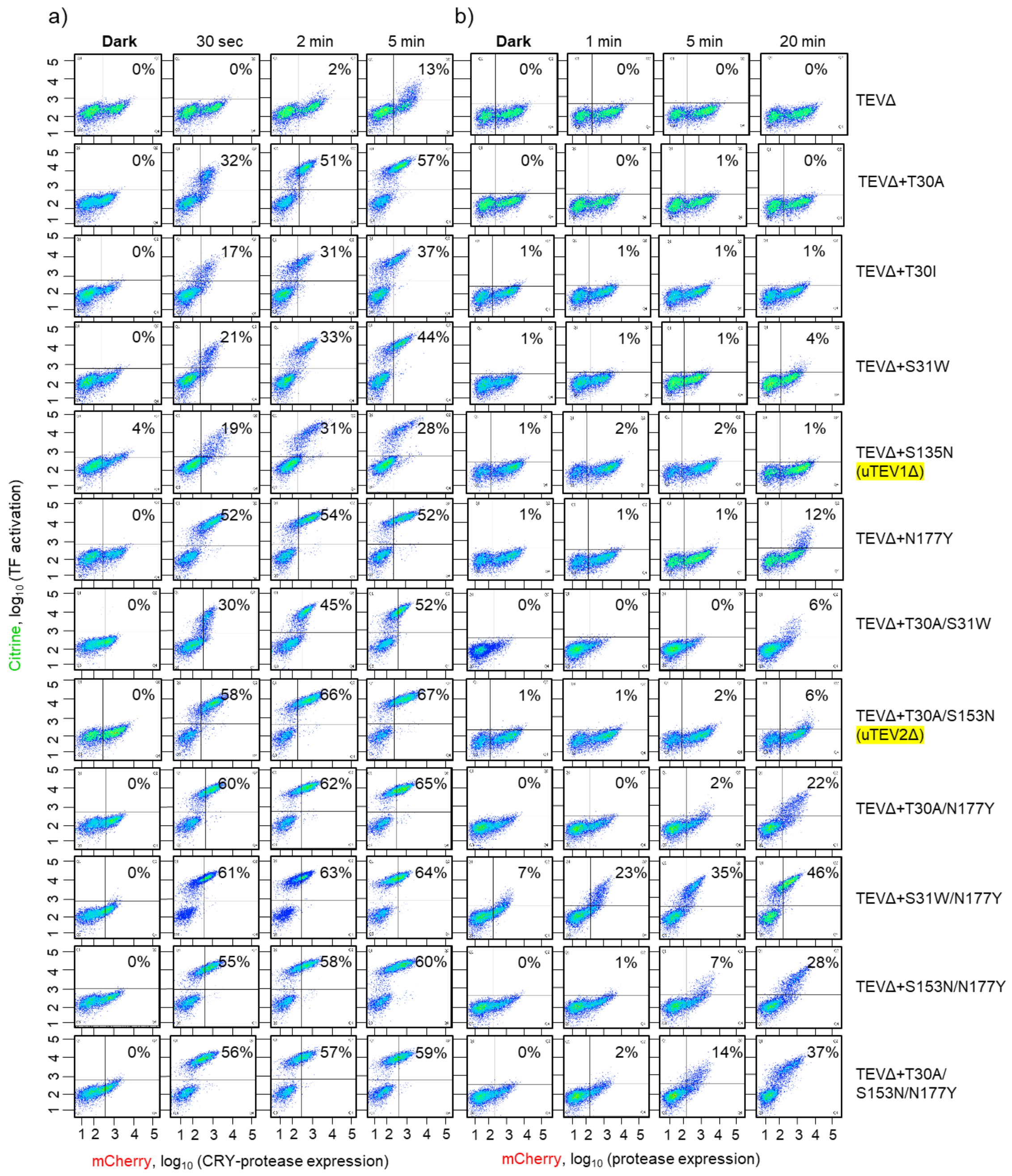


**Supplementary Figure 3. Characterization of evolved TEVΔ mutants in yeast.** Related to Figure 2A. (**A**) FACS plots were collected 6 hours after blue light exposure for the indicated times (0.5, 2, and 5 min). Percentages of Citrine-positive cells are shown in the graph in Figure 2A. (**B**) Same as (A) but with CRY omitted to test for proximity-dependence of TEVΔ-TEVcs interaction (cells express TEVΔ-mCherry instead of CRY-TEVΔ-mCherry). Each plot represents two replicates, n = 10,000 cells.

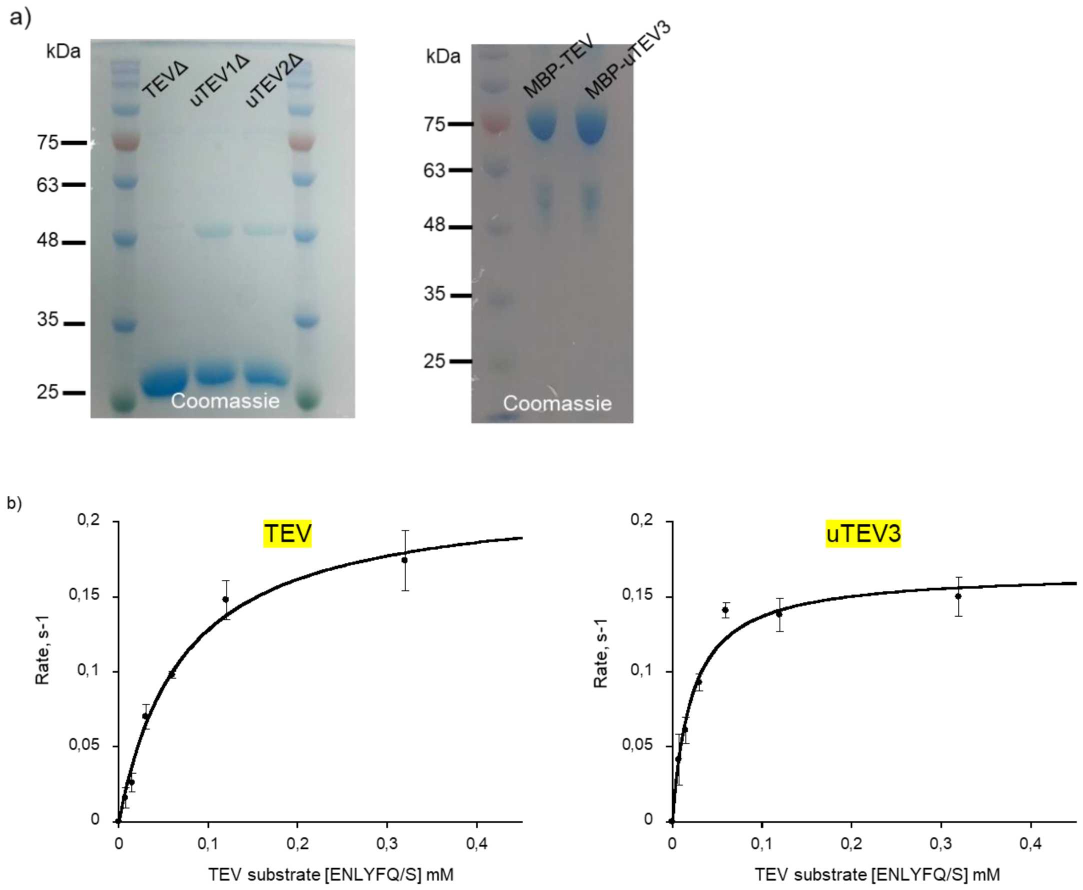


**Supplementary Figure 4**. **TEV purification and kinetics**. Related to Figures 2C-D and 3E-F. (**A**) SDS-PAGE (9%) of purified TEV proteases. TEVΔ is 25 kD. MBP-TEV (full-length) is 75 kD. (**B**) Michaelis-Menten plots for wild-type full-length TEV and uTEV3 (containing mutations I138T, S135N, and T180A). Reactions were assembled with 100 nM purified protease and variable amounts (0.0075-0.32 mM) of substrate protein MBP-TEVcs(ENLYFGS)-GFP at 30 °C. Initial proteolysis rates were determined for each starting substrate concentration, using the in-gel fluorescence assay shown in Figures 2C and 3E. Data was fit to a Michaelis-Menten enzyme kinetics model with center values representing the mean and error bars representing the standard deviation of three technical replicates.

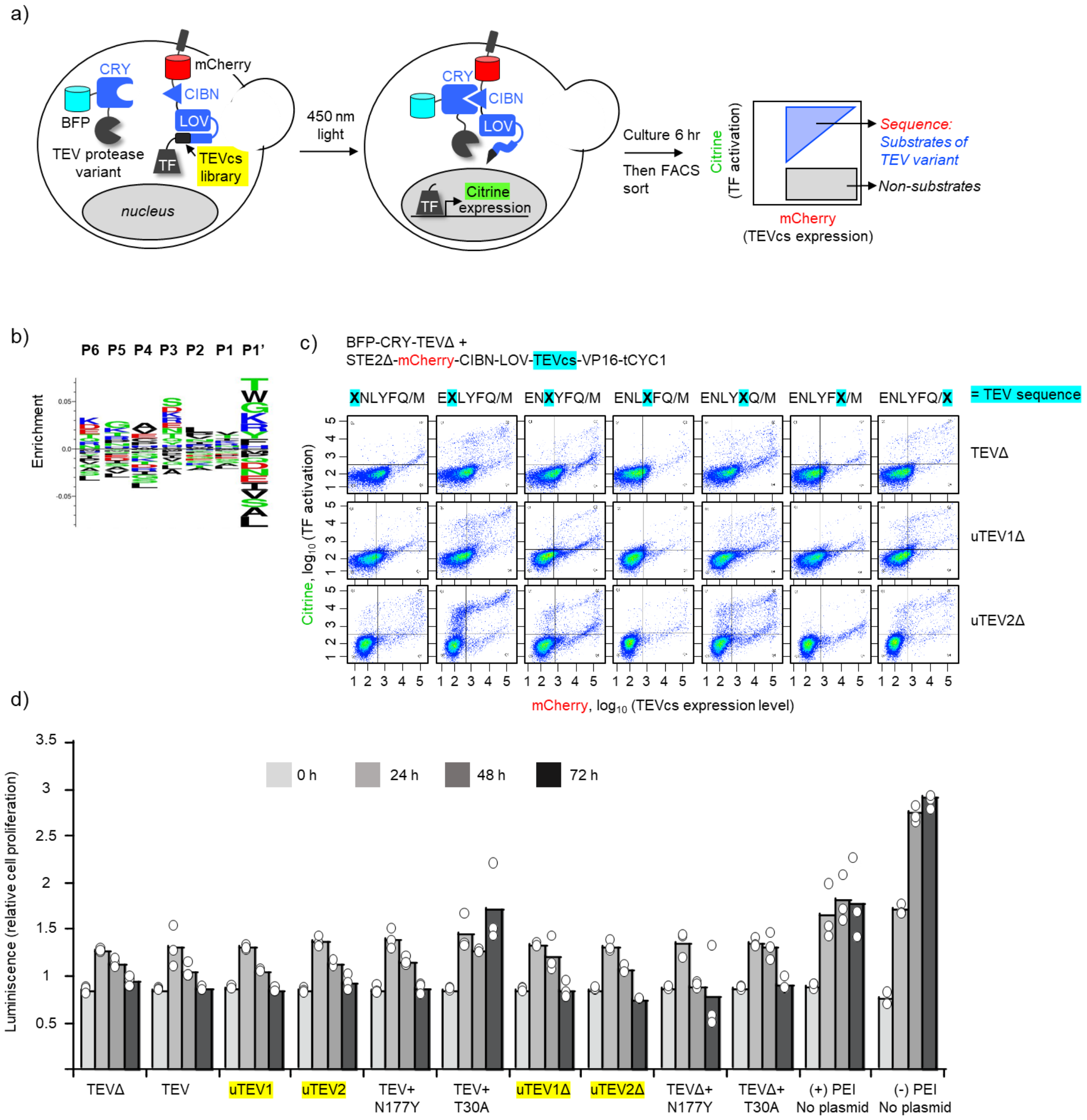


**Supplementary Figure 5. Profiling the sequence specificity of TEVΔ variants in yeast**. Related to Figure 2F-G**.** (**A**) Assay for profiling yeast sequence specificity. The protease variant of interest is co-expressed with a library of TEVcs sequences (flanked by LexA-VP16 TF at the C-terminal end, and a plasma membrane anchor, mCherry, CIBN, and LOV at the N-terminal end). Upon exposure of cells to blue 450 nm light, the CRY-CIBN interaction brings the protease proximal to TEVcs, and the LOV domain changes conformation to expose TEVcs. Sequences sensitive to TEV proteolysis will release the TF, which translocates to the nucleus and drives expression of the reporter gene Citrine. (**B**) Sequence profile of the seven TEVcs libraries with randomized nucleotides (pooled together) before sorting. Each of the seven TEVcs libraries is randomized at a single position only. (**C**) Analysis of single randomized positions in the TEV cleavage site, using wild-type TEVΔ, uTEV1Δ, and uTEV2Δ. Sample FACS plots 6 hours after blue light exposure. Each plot represents one replicate, n = 10,000 cells. (**D**) Viability assays in HEK 293T expressing the evolved TEV proteases. This experiment was performed once, with three biological replicates per sample. White dots indicate individual technical replicates.

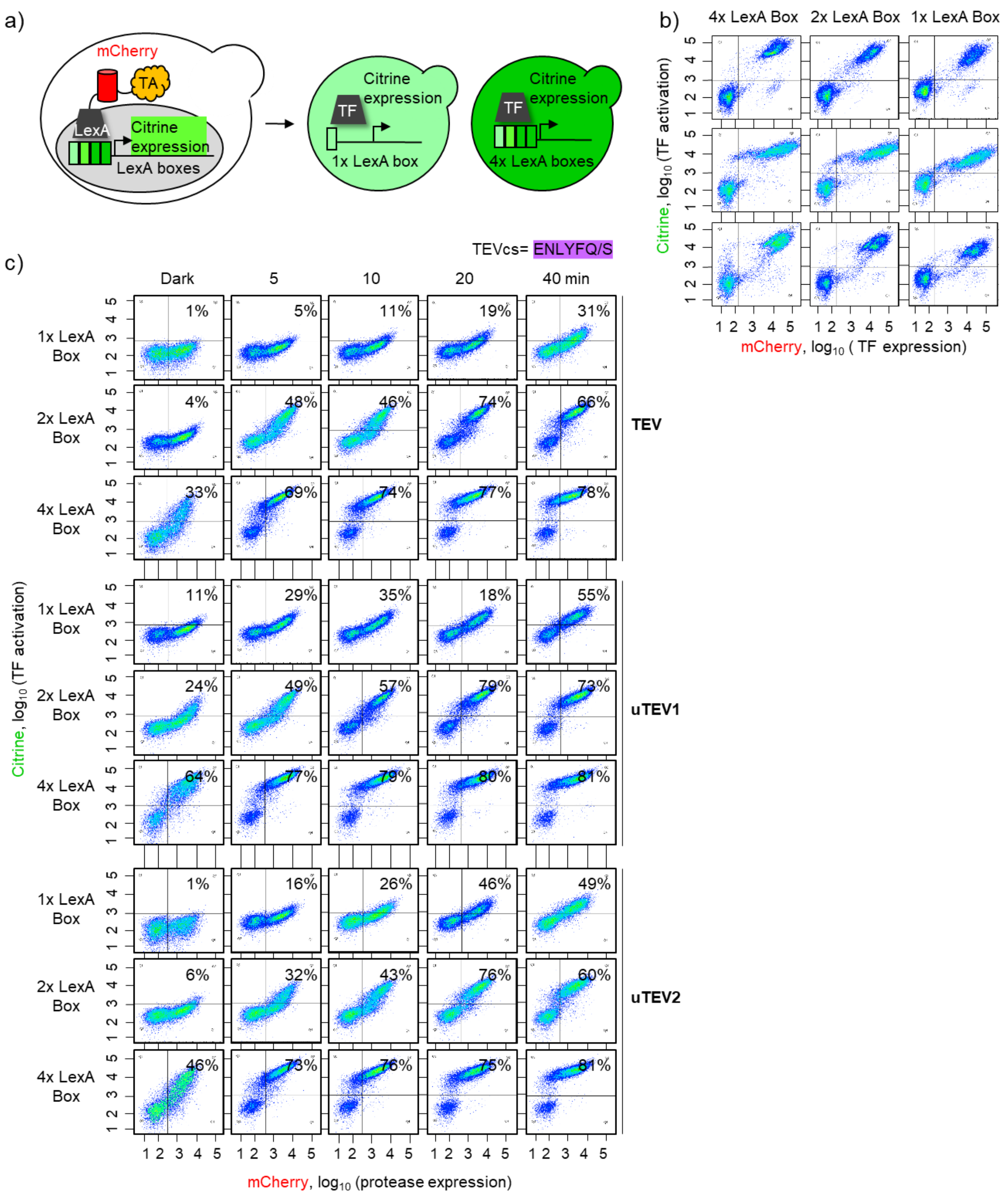


**Supplementary Figure 6. Optimizing the yeast platform for evolution of full-length high-affinity proteases.** Related to Figure 3. (**A**) To tune the dynamic range of the platform, the LexA DNA-binding domain was fused to one of 3 different transcriptional activators (TAs): VP16, B42, or Gal4. The constructs were expressed in yeast containing different numbers of LexA boxes upstream of the Citrine reporter gene. (**B**) FACS data showing the effect of varying the number of LexA boxes in the promoter with different TAs. FACS data collected 12 hours following galactose induction. Each plot represents two replicates, n = 20,000 cells. (C) Comparison of different full-length TEVs in the setup shown in Figure 3A. The TEVcs was the high-affinity sequence ENLYFQ/S. FACS plots obtained 6 hours after blue light exposure for the indicated times (5, 10, 20 and 40 min). Each plot represents one replicate, n = 20,000 cells. Percentage of Citrine-positive cells in each condition used to generate the graph in Figure 3B.

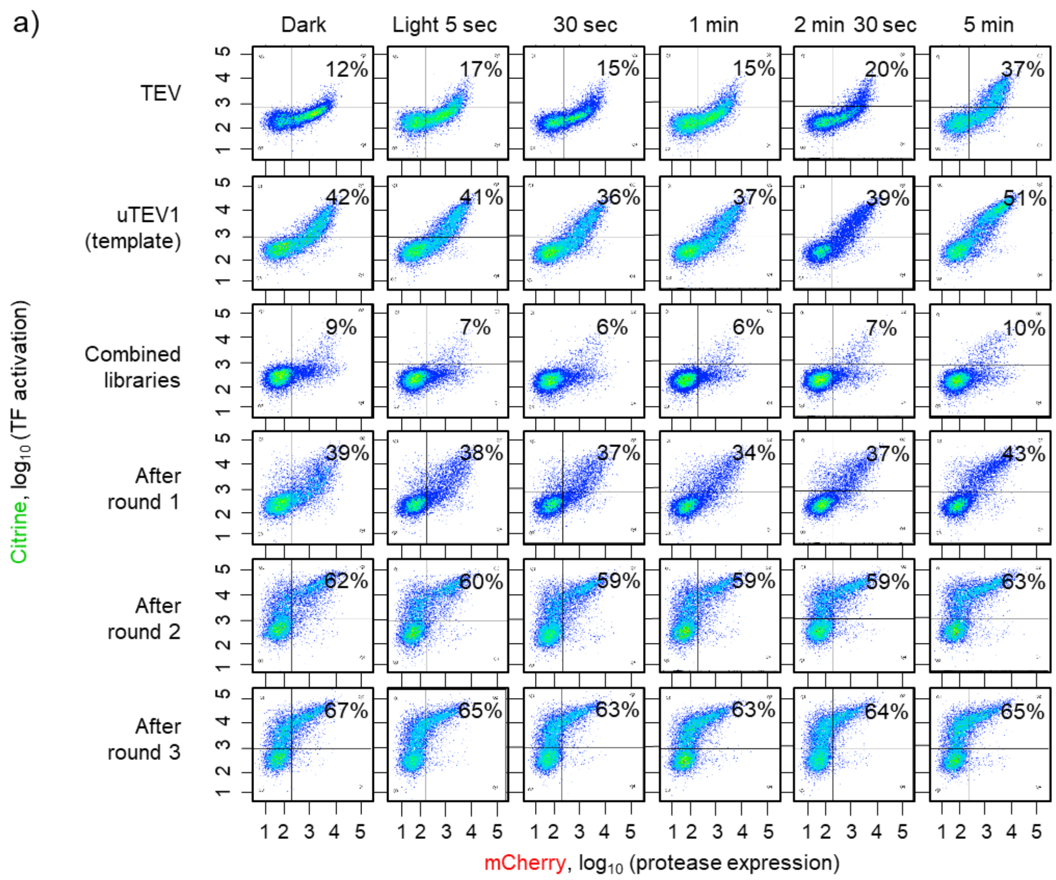


b)

| Clone | Mutations |
|---|---|
| #1 | S3P, T180A |
| #2 | N177Y, F225L |
| #3 | S135F |
| #4 | T30I, D90G, T158A, E237G |
| #5 | K119E, T180A, N236S |
| #6 | template |
| #7 | N68D, P93S, S122P, S135F |
| #8 | K67E, S135F, M218T |
| #9 | M87L, T180A |
| #10 | M87T, I138T |
| #11 | F132S, N153D |
| #12 | T180A |
| #13 | K141R |
| #14 | S135F, K147E |
| #15 | template |
| #16 | S122P, T180A |
| #17 | P224S, N236S |
| #18 | N205D |
| #19 | E106G, I138T |
| #20 | T180A |
| #21 | inserted linker (GSASGSGKLSGGG), D90G, I138T, K220R |
| #22 | empty vector |
| #23 | T180A |
| #24 | S135F |

**Supplementary Figure 7. Analysis of selected yeast populations and full-length TEV clones.** Related to Figure 3. (**A**) Same as Figure 3C but showing more conditions. (**B**) Sequencing after round 3. 24 clones were sequenced and mutations found in each, relative to the original template uTEV1, are shown.

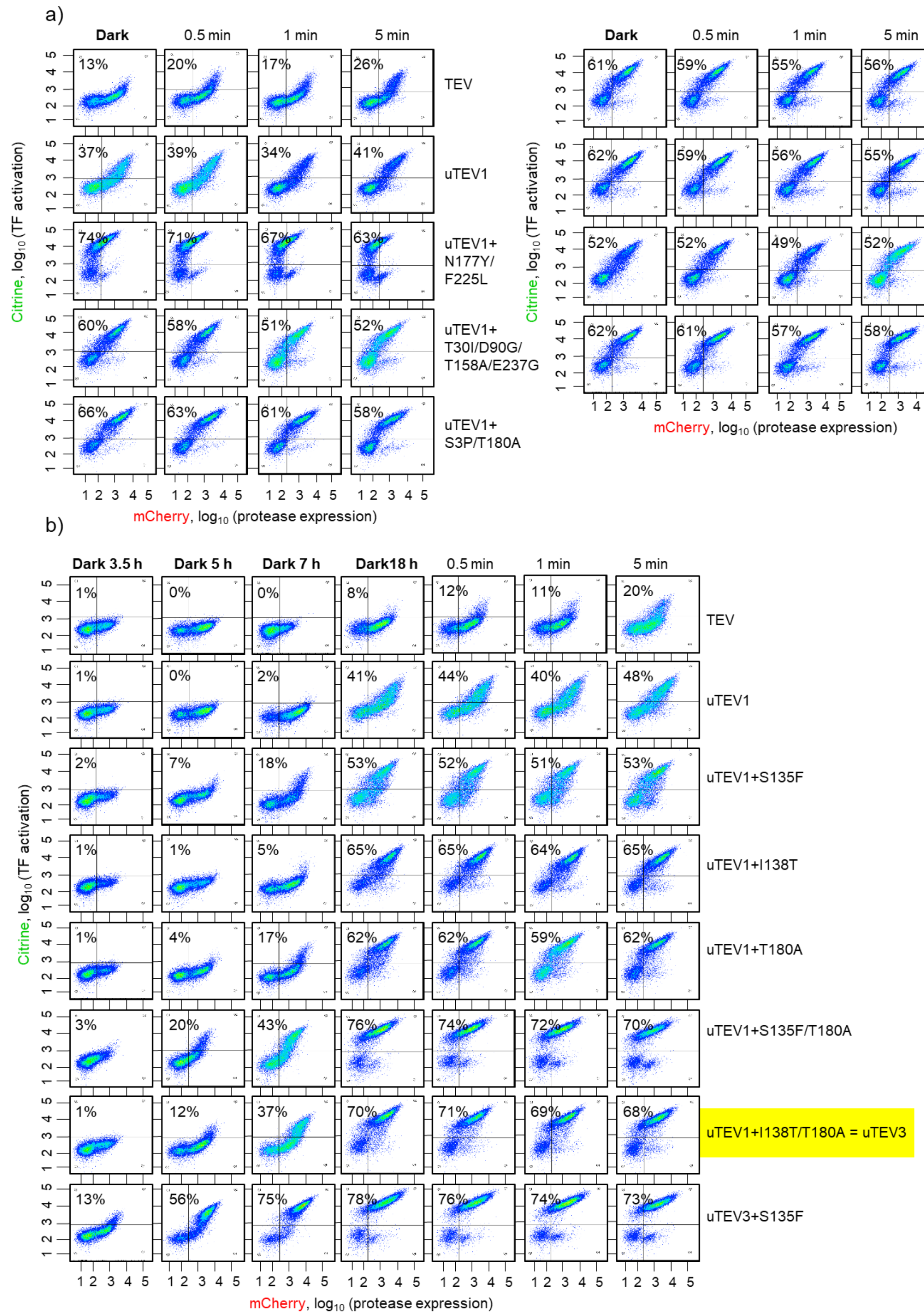


**Supplementary Figure 8. Characterization of evolved full-length TEV mutants.** Related to Figure 3D. (**A**) Proteases were expressed in a yeast strain with 2 LexA boxes and high-affinity TEVcs (ENLYFQ/S) (configuration shown in Figure. 3A). FACS analysis performed 6 hours after light irradiation. Each condition repeated once. n = 20,000 cells. (**B**) Same as (A) with additional TEV mutants and additional conditions. The first 3 columns show shorter protein induction times in the dark (standard induction time is 12 hours). Right 3 columns show cells 6 hours following blue light irradiation for the indicated times. Each condition performed twice. n = 20,000 cells.

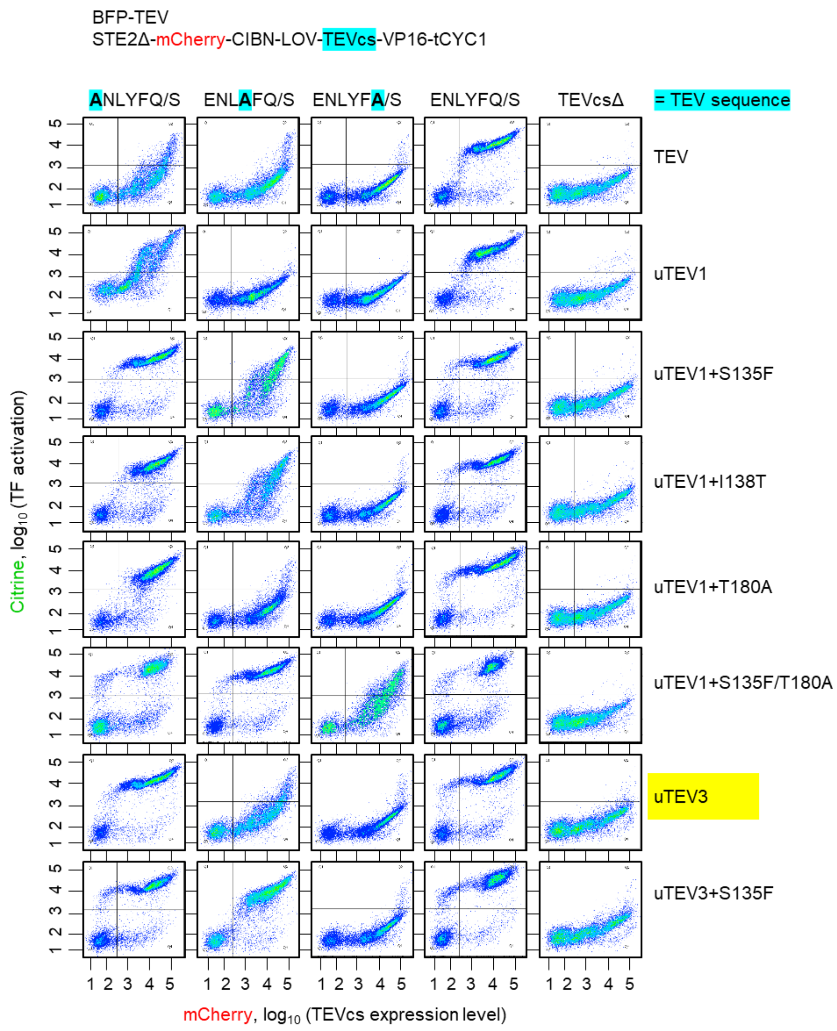


**Supplementary Figure 9. Profiling the sequence specificity of full-length TEV mutants in yeast**. Same assay as in Figure 2F. FACS analysis performed 12 hours after galactose induction. Each condition performed once, n = 20,000 cells.

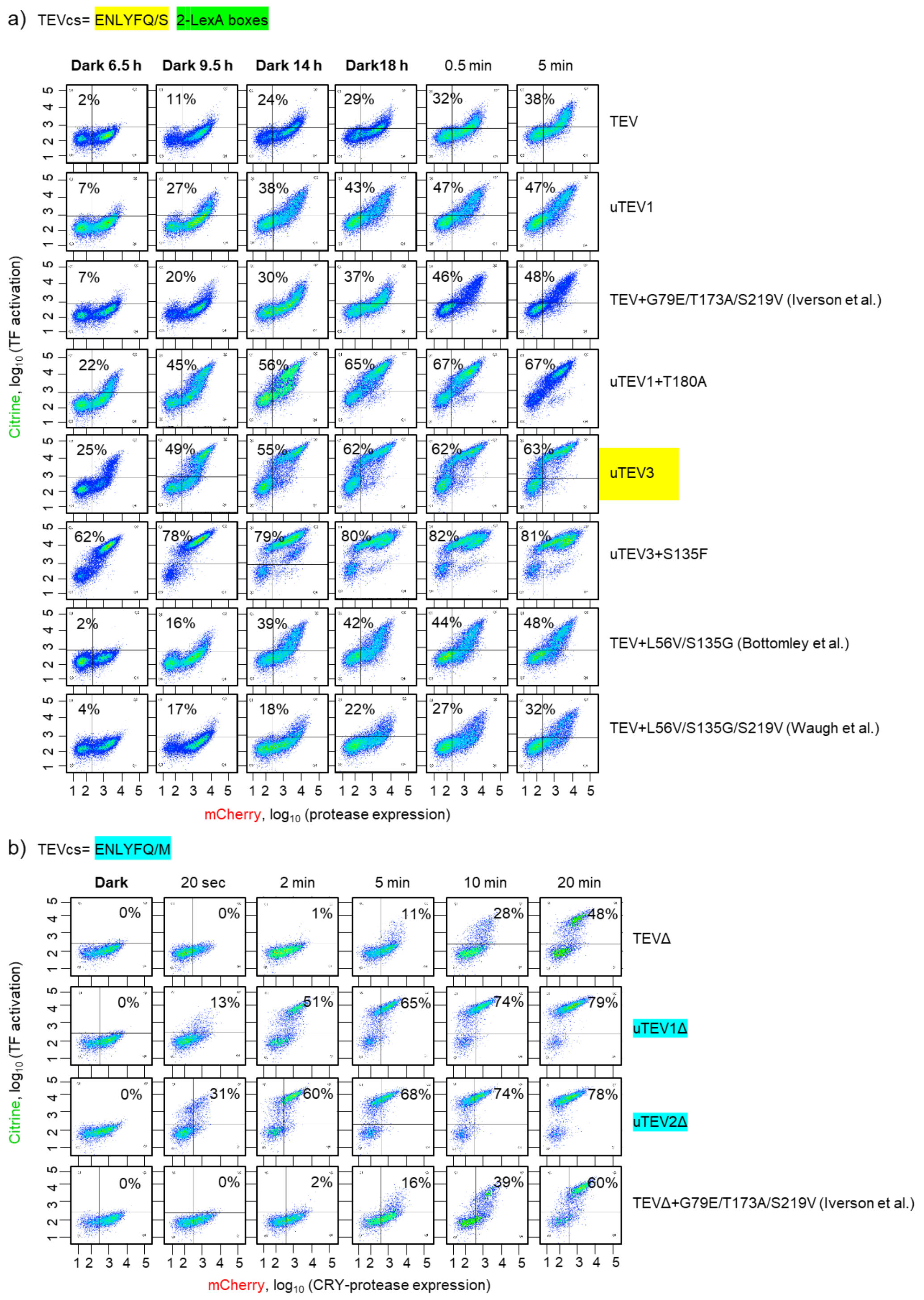


**Supplementary Figure 10. Comparison of evolved TEVs with Iverson, Bottomley and Waugh TEV mutants.** (**A**) Side-by-side comparison in yeast, with full-length proteases and the high-affinity TEVcs (ENLYFQ/S). First four columns show yeast induced with galactose in the dark for 6.5 to 18 hours before FACS analysis. Last two columns were irradiated with light before FACS analysis 6 hours later. (**B**) Side-by-side comparison of truncated proteases using the low-affinity TEVcs (ENLYFQ/M). FACS analysis was performed 6 hours after blue light exposure for the indicated times. Each condition was repeated once, n = 20,000.

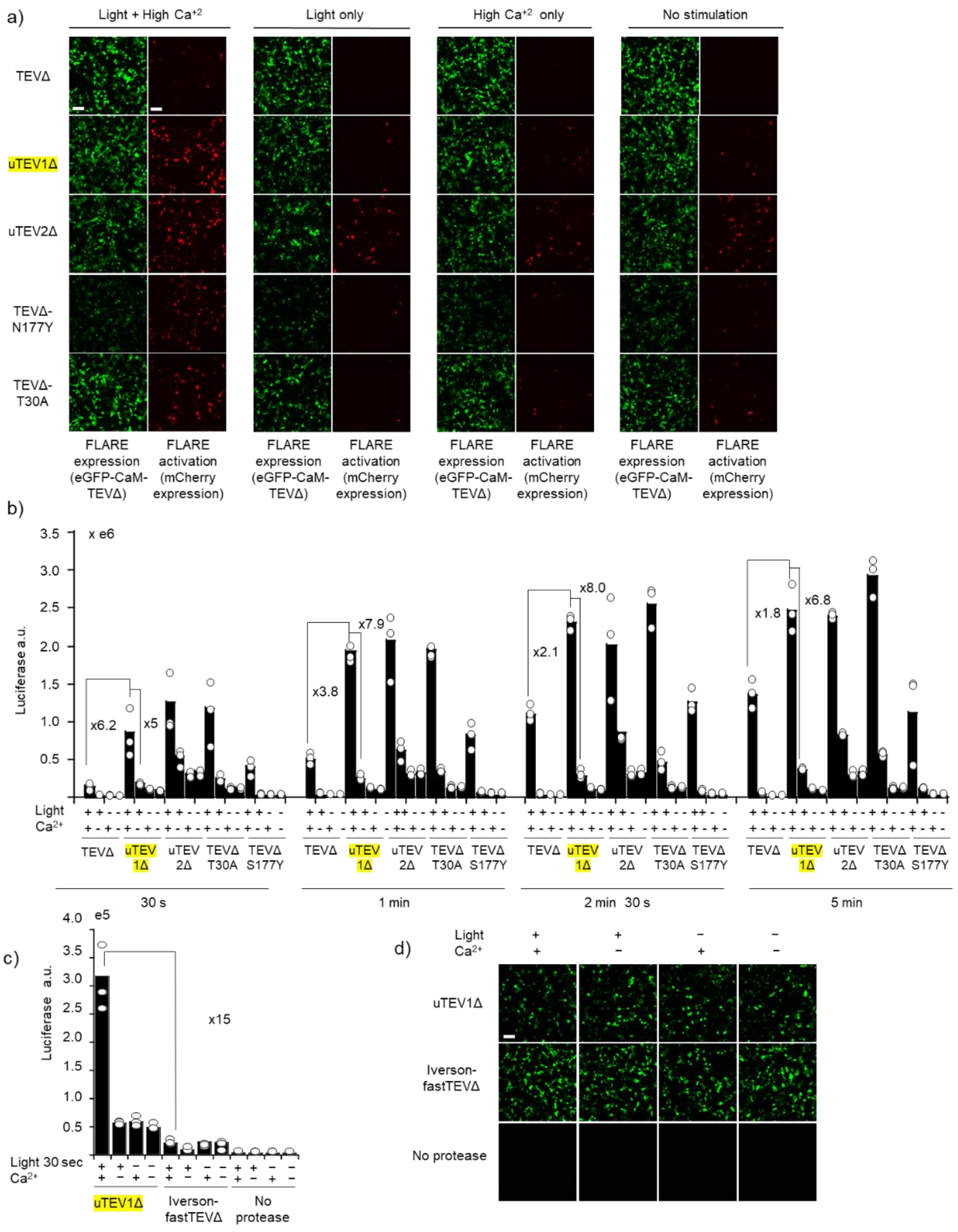


**Supplementary Figure 11**. **Testing evolved TEV mutants in FLARE**. Related to Figure 4E. (**A**) HEK293T cells were transiently transfected with FLARE constructs (as in Figure 4A) incorporating the indicated TEV protease. Stimulation was performed using 5 mM $CaCl_2$ and ionomycin for 30 seconds in the presence of blue light (467 nm, 60 mW/cm2, 10% duty cycle (0.5s light every 5s). Nine hours later, cells were fixed and imaged. This experiment was performed independently two times with similar results. (**B**) Same experiment as in (A) but with luciferase as the reporter gene instead of mCherry. Stimulation times varied from 30 seconds to 5 minutes. This experiment was performed once with three technical replicates per condition. (**C**) Comparison of uTEV1Δ with the truncated version of Iverson's TEV in the context of FLARE. Cells were stimulated and analyzed as in (B). This experiment was performed once, with three technical replicates per condition. (**D**) Samples from (C) were imaged by confocal microscopy to confirm protease expression. GFP channel shown. Scale bar, 10 µm.

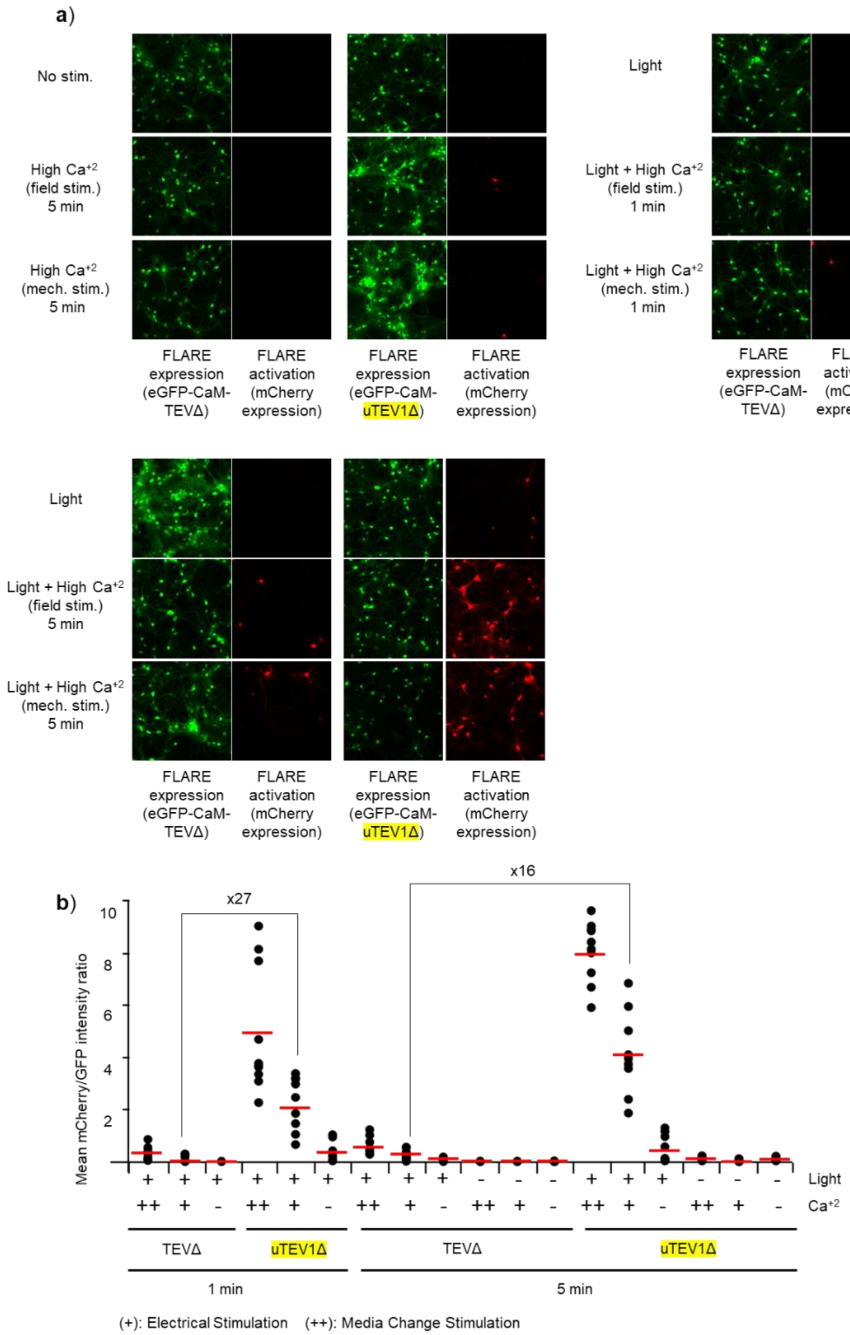


**Supplementary Figure 12**. **Evaluation of uTEV1Δ in the context of FLARE in neurons**. Related to Figure 4F. (**A**) Rat cortical neurons were transduced at day 12 with FLARE constructs (packaged into AAV1/2 viruses) containing either the original TEVΔ protease or our evolved uTEV1Δ protease. At day 18 in vitro (DIV18), we stimulated the neurons using either field stimulation (3-s trains consisting of 32 1-ms 50 mA pulses at 20 Hz for a total of 1 or 5 min), or via replacement of culture media with media of identical composition (this mechanically stimulates the cultures and also provides a fresh source of glutamate). Light source was 467 nm, 60 mW/cm2, 10% duty cycle (0.5s light every 5s). Imaging was performed 18 hours later. This experiment was replicated three times for each condition. Scale bars, 10 µm. (**B**) Quantitation of data from (A). Signal ratios were calculated from mean mCherry and mean GFP intensities across >50 cells per field of view. 10 fields of view quantified per condition. Red lines indicates the mean of 10 FOVs.

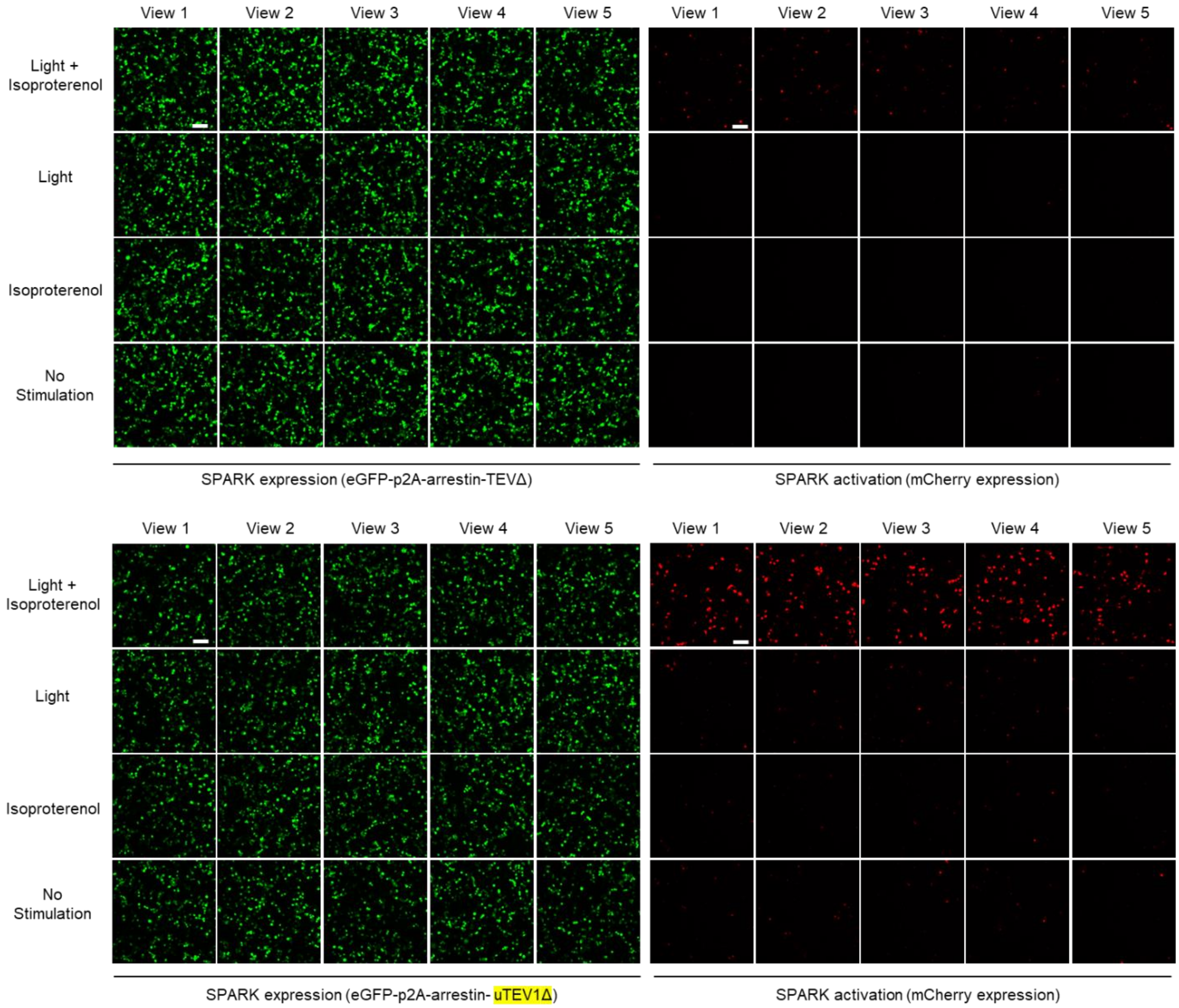


**Supplementary Figure 13**. **Testing the evolved protease uTEV1Δ in SPARK**. Related to Figure 4G. HEK293T cells were transiently transfected with SPARK constructs (Figure 4B) containing the indicated protease variant. Cells were stimulated with 10 μM isoproterenol for 60 sec in the presence or absence of blue light (467 nm, 60 mW/cm2, 10% duty cycle (0.5s light every 5s)). Nine hours later, cells were imaged. This experiment was replicated two times. Scale bars, 10 μm.

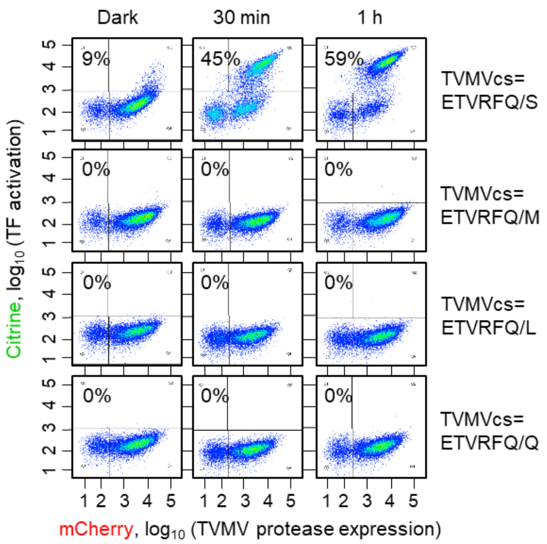


**Supplementary Figure 14. Testing a different protease (TVMV) in the yeast platform.** Constructs were designed as in Figure 1A, but TEV was replaced with TVMV protease, and TEVcs was replaced with the TVMV substrate sequences shown at right. FACS plots were collected 6 hours after blue light irradiation for the indicated times. Percentages give the fraction of Citrine-positive cells. Each plot representative of two replicates, n = 20,000 cells.

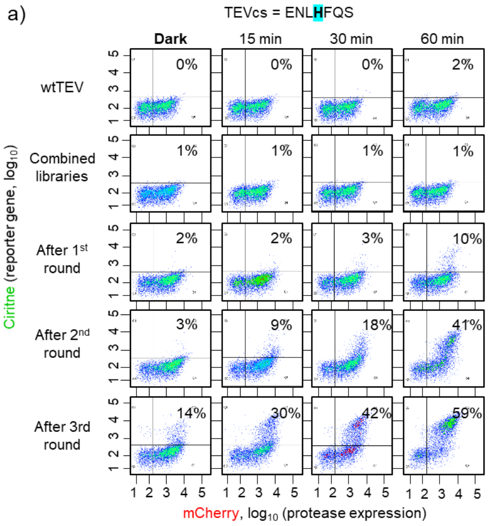


C) TEVcs = ENLHFQ/S

| | | | | | | | | | | | | | | | |
|---|---|---|---|---|---|---|---|---|---|---|---|---|---|---|---|
| #1 | | | | | | | | | | | | N177Y | | | |
| #2 | | | | | K97R | | | | I138T | | | N177Y | | | V239A |
| #3 | | | | | | | | | I138T | | | N177Y | | | |
| #4 | | | | D90G | | | | | | | | N177Y | | | |
| #5 | | | | | | | | | | Y140H | K141R | N177Y | | | |
| #6 | | | | | | | | D127 G | | | | N177Y | | | |
| #7 | Template | | | | | | | | | | | | | | |
| #8 | | | | | K97R | | | | I138T | | | N177Y | | | V239A |
| #9 | | | | | K97R | | | | I138T | | | N177Y | | | V239A |
| #10 | | | | | | | | | | | | N177Y | | | |
| #11 | | | | | | | F125L | | | | | N177Y | | | |
| #12 | | | | | k97R | | | | I138T | | | N177Y | | | V239A |
| #13 | | | | | | | | D127 G | | | | N177Y | | | |
| #14 | | | | | | | | | | | | N177Y | | | |
| #15 | | | | | | | | | | | K141R | N177Y | T180A | | |
| #16 | | | M87T | | | | | | | | | N177Y | | | |
| #17 | | | | | | | | | I138T | | | N177Y | | | |
| #18 | | | | | | | | | | | | N177Y | T180A | | |
| #19 | | | | | | | | | | | | N177Y | | | |
| #20 | | H28Y | | | | | | | | | | N177Y | | | |
| #21 | | | | | | | | | | | K141R | N177Y | T180A | | |
| #22 | | | | | | | | | | | | N177Y | | | |
| #23 | | | | | | P103L | | | | | | N177Y | | M235T | |
| #24 | | | | | | | | | | | K141R | N177Y | T180A | | |

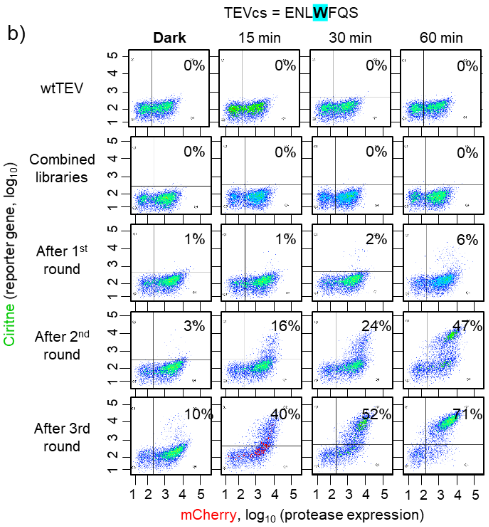


d) TEVcs = ENLWFQ/S

| | | | | | | | | | | | | | | | | |
|---|---|---|---|---|---|---|---|---|---|---|---|---|---|---|---|---|
| #1 | | | | | | | N177Y | | N192D | | | | | | | |
| #2 | | | | | | | N177Y | | | | | | | | | |
| #3 | | | | | | | N177Y | | | | | | | | | |
| #4 | | | | | | | N177Y | | | | | | | | | |
| #5 | | | | | | F132S | N177Y | | | | | | | | | |
| #6 | | | | | | | N177Y | | | N215D | | | | | | |
| #7 | | | | | | | N177Y | | | | | | | | | |
| #8 | | | | | | | N177Y | | | | | | | | | |
| #9 | | | | | | | N177Y | | | | | | | | | |
| #10 | | | | | | | N177Y | | | | S219G | | | | | |
| #11 | | | | | | | N177Y | | | | | | | | | |
| #12 | | | | | | | N177Y | | | | | | | P224L | | |
| #13 | | | F40S | | | | N177Y | | | | | | | | | |
| #14 | Empty Vector | | | | | | | | | | | | | | | |
| #15 | | | | | | | N177Y | | | | | | | | | |
| #16 | | | | Q117 R | | | N177Y | | | | | | | | | |
| #17 | | | | | | | N177Y | | | | | | | | | Y240D |
| #18 | | | | | | | N177Y | | | | | | | | | |
| #19 | | | | | | | N177Y | | | | | | | | | |
| #20 | | | | | S120G | | N177Y | K184R | | | | | | | | |
| #21 | | S16N | | | | | N177Y | | | | | | | | | |
| #22 | | | | | | | N177Y | | | | | P221S | | | L238S | |
| #23 | | | | | | | N177Y | | | | | | | | | Y240N |
| #24 | | | | | | | N177Y | | | | | | E222G | | | |

**Supplementary Figure 15. Selections to alter the sequence-specificity of TEV.** Analysis of full-length TEV libraries after 3 rounds of sorting against mutated TEV cleavage sequences: H at the P3 position of TEVcs in (**A**), and W in the P3 position of TEVcs in (**B**). The configuration of constructs was the same as in Figure 3A. FACS plots were obtained 6 hours after blue light exposure for the indicated times. Percentages quantify Citrine-positive cells. This experiment was performed once, n = 10,000 cells. (**C**) and (**D**) Sequencing after round 3. 24 clones were sequenced from each selection and mutations found in each, relative to the original template wild-type TEV, are shown.

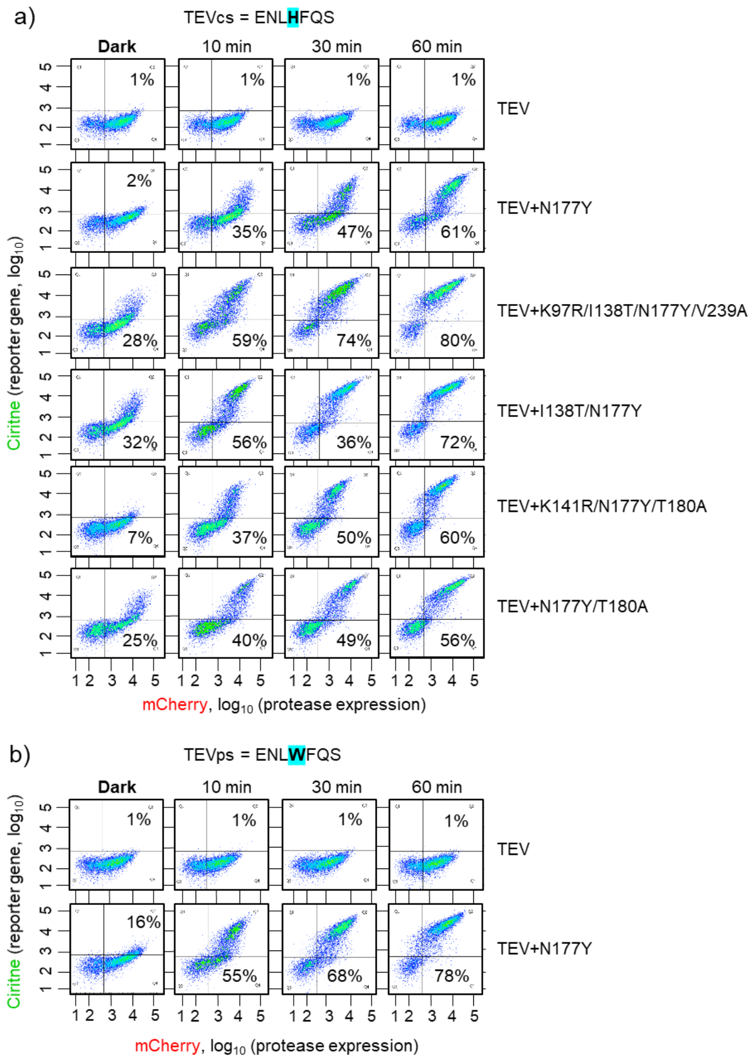


**Supplementary Figure 16. Characterization of specificity-altered TEV mutants in yeast.** Related to Supplementary Figure 15. The indicated TEV mutants were compared to wild-type full-length TEV, using the altered TEVcs substrates P3 = H (**A**) and P3 = W (**B**). FACS plots obtained 6 hours after blue light exposure for the indicated times. Percentages reflect the fraction of Citrine-positive cells**.** Each plot representative of two replicates, n = 10,000 cells.

### Plasmid table

| ID | Plasmid name | Plasmid Vect | Promoter | Terminator | Expression in | Tags | More Details | Ab. R | Used in figures |
|---|---|---|---|---|---|---|---|---|---|
| p1 | STE2-V5-CIBN-M13-eLOV-TEVcs(ENLYFQM)-LexAVP16 | pRS-derived | ACT1 | | Yeast / ApaI digestion to integrate | V5 | HIS3 selectable marker | Amp | Fig. 1 / S. Fig 1B |
| p2 | STE2-V5-BFP-CIBN-M13-eLOV-TEVcs(ENLYFQM)-LexAVP16 | pRS-derived | ACT1 | | Yeast / ApaI digestion to integrate | V5 / BFP | HIS3 selectable marker | Amp | Fig. 1 / S. Fig 1B |
| p3 | STE2Δ-BFP-CIBN-MKII-eLOV-TEVcs(ENLYFQM)-LexAVP16-tC | pRS-derived | TDH3 | tCYC1 | Yeast / AscI digestion to integrate | flag / BFP | LEU2 selectable marker | Amp | Fig. 1 / S. Fig 1B |
| p4 | STE2Δ-CIBN-MKII-eLOV-TEVcs(ENLYFQM)-LexAVP16-tCYC | pRS-derived | TDH3 | tCYC1 | Yeast / AscI digestion to integrate | flag / BFP | LEU2 selectable marker | Amp | Fig. 1 / S. Fig 1B |
| p5 | STE2Δ-V5-CIBN-M13-eLOV(ENLYFQM)-TEVcs-LexAVP16 | pRS-derived | ACT1 | | Yeast / AscI digestion to integrate | V5 | LEU2 selectable marker | Amp | Fig. 1 / S. Fig 1B |
| p6 | STE2Δ-V5-BFP-CIBN-M13-eLOV(ENLYFQM)-TEVcs-LexAVP1( | pRS-derived | ACT1 | | Yeast / AscI digestion to integrate | V5 / BFP | LEU2 selectable marker | Amp | Fig. 1 / S. Fig 1B |
| p7 | mCherry-CRY-TEV | pRSII415 | Gal | tCYC1 | Yeast / episomally | mCherry | LEU2 selectable marker | Amp | Fig. 1 / S. Fig 1B |
| p8 | mCherry-CRY-TEVΔ/S219V | pRSII415 | Gal | tCYC1 | Yeast / episomally | mCherry | LEU2 selectable marker | Amp | Fig. 1 / S. Fig 1B |
| p9 | mCherry-TEV | pRSII415 | Gal | tCYC1 | Yeast / episomally | mCherry | LEU2 selectable marker | Amp | Fig. 1 / S. Fig 1B |
| p10 | mCherry-TEVΔ-S219V | pRSII415 | Gal | tCYC1 | Yeast / episomally | mCherry | LEU2 selectable marker | Amp | Fig. 1 / S. Fig 1B |
| p11 | mCherry-CRY-TEV | pRSII413 | Gal | tCYC1 | Yeast / episomally | mCherry | HIS3 selectable marker | Amp | Fig. 1 / S. Fig 1B |
| p12 | mCherry-CRY-TEVΔ/S219V | pRSII413 | Gal | tCYC1 | Yeast / episomally | mCherry | HIS3 selectable marker | Amp | Fig. 1 / S. Fig 1B |
| p13 | mCherry-TEV | pRSII413 | Gal | tCYC1 | Yeast / episomally | mCherry | HIS3 selectable marker | Amp | Fig. 1 / S. Fig 1B/ |
| | | | | | | | | | Fig. 2 S. Fig. 3 |
| p14 | mCherry-TEVΔ/S219V | pRSII413 | Gal | tCYC1 | Yeast / episomally | mCherry | HIS3 selectable marker | Amp | Fig. 1 / S. Fig 1B/ Fig. 2 |
| | | | | | | | | | S. Fig. 3 / S. Fig 10 |
| p15 | STE2Δ-BFP-CIBN-MKII-eLOV-TEVcs-LexAB42 | pRS-derived | ACT1 | tCYC1 | Yeast / AscI digestion to integrate | V5 / BFP | LEU2 selectable marker | Amp | S. Fig 1C |
| p16 | STE2Δ-BFP-CIBN-MKII-eLOV-TEVcs-LexAGal4 | pRS-derived | ACT1 | tCYC1 | Yeast / AscI digestion to integrate | V5 / BFP | LEU2 selectable marker | Amp | S. Fig 1C |
| p17 | mCherry-CRY-TEVΔ/S219V+S31W | pRSII413 | Gal | tCYC1 | Yeast / episomally | mCherry | HIS3 selectable marker | Amp | Fig. 2 / S. Fig 3 |
| p18 | mCherry-CRY-TEVΔ/S219V+S153N | pRSII413 | Gal | tCYC1 | Yeast / episomally | mCherry | HIS3 selectable marker | Amp | Fig. 2 / S. Fig 3 / S. Fig 10 |
| p19 | mCherry-CRY-TEVΔ/S219V+T30A+S153N | pRSII413 | Gal | tCYC1 | Yeast / episomally | mCherry | HIS3 selectable marker | Amp | Fig. 2 / S. Fig 3 |
| p20 | mCherry-CRY-TEVΔ/S219V+T30I | pRSII413 | Gal | tCYC1 | Yeast / episomally | mCherry | HIS3 selectable marker | Amp | Fig. 2 / S. Fig 3 |
| p21 | mCherry-CRY-TEVΔ/S219V+N177Y | pRSII413 | Gal | tCYC1 | Yeast / episomally | mCherry | HIS3 selectable marker | Amp | Fig. 2 / S. Fig 3 |
| p22 | mCherry-CRY-TEVΔ/S219V+T30A | pRSII413 | Gal | tCYC1 | Yeast / episomally | mCherry | HIS3 selectable marker | Amp | Fig. 2 / S. Fig 3 |
| p23 | mCherry-TEVΔ/S219V+S31W | pRSII413 | Gal | tCYC1 | Yeast / episomally | mCherry | HIS3 selectable marker | Amp | Fig. 2 / S. Fig 3 |
| p24 | mCherry-TEVΔ/S219V+S153N | pRSII413 | Gal | tCYC1 | Yeast / episomally | mCherry | HIS3 selectable marker | Amp | Fig. 2 / S. Fig 3 |
| p25 | mCherry-TEVΔ/S219V+T30A+S153N | pRSII413 | Gal | tCYC1 | Yeast / episomally | mCherry | HIS3 selectable marker | Amp | Fig. 2 / S. Fig 3 |
| p26 | mCherry-TEVΔ/S219V+T30I | pRSII413 | Gal | tCYC1 | Yeast / episomally | mCherry | HIS3 selectable marker | Amp | Fig. 2 / S. Fig 3 |
| p27 | mCherry-TEVΔ/S219V+N177Y | pRSII413 | Gal | tCYC1 | Yeast / episomally | mCherry | HIS3 selectable marker | Amp | Fig. 2 / S. Fig 3 |
| p28 | mCherry-TEVΔ/S219V+T30A | pRSII413 | Gal | tCYC1 | Yeast / episomally | mCherry | HIS3 selectable marker | Amp | Fig. 2 / S. Fig 3 |
| p29 | mCherry-CRY-TEVΔ/S219V+T30A+N177Y | pRSII413 | Gal | tCYC1 | Yeast / episomally | mCherry | HIS3 selectable marker | Amp | Fig. 2 / S. Fig 3 |
| p30 | mCherry-CRY-TEVΔ/S219V+S153N+N177Y | pRSII413 | Gal | tCYC1 | Yeast / episomally | mCherry | HIS3 selectable marker | Amp | Fig. 2 / S. Fig 3 |
| p31 | mCherry-CRY-TEVΔ/S219V+S31W+N177Y | pRSII413 | Gal | tCYC1 | Yeast / episomally | mCherry | HIS3 selectable marker | Amp | Fig. 2 / S. Fig 3 |
| p32 | mCherry-CRY-TEVΔ-S219V- T30A- S153N + Asn177Tyr-tCYC1 | pRSII413 | Gal | tCYC1 | Yeast / episomally | mCherry | HIS3 selectable marker | Amp | Fig. 2 / S. Fig 3 |
| p33 | mCherry-CRY-TEVΔ-S219V- Thr30Ala + Ser31Trp-tCYC1 | pRSII413 | Gal | tCYC1 | Yeast / episomally | mCherry | HIS3 selectable marker | Amp | Fig. 2 / S. Fig 3 |
| p34 | mCherry-TEVΔ/S219V+T30A+N177Y | pRSII413 | Gal | tCYC1 | Yeast / episomally | mCherry | HIS3 selectable marker | Amp | Fig. 2 / S. Fig 3 |
| p35 | mCherry-TEVΔ/S219V+S153N+N177Y | pRSII413 | Gal | tCYC1 | Yeast / episomally | mCherry | HIS3 selectable marker | Amp | Fig. 2 / S. Fig 3 |
| p36 | mCherry-TEVΔ/S219V+S31W+N177Y | pRSII413 | Gal | tCYC1 | Yeast / episomally | mCherry | HIS3 selectable marker | Amp | Fig. 2 / S. Fig 3 |
| p37 | mCherry-TEVΔ/S219V+T30A+S153N+N177Y | pRSII413 | Gal | tCYC1 | Yeast / episomally | mCherry | HIS3 selectable marker | Amp | Fig. 2 / S. Fig 3 |
| p38 | mCherry-TEVΔ/S219V+T30A+S31W | pRSII413 | Gal | tCYC1 | Yeast / episomally | mCherry | HIS3 selectable marker | Amp | Fig. 2 / S. Fig 3 |
| p39 | mCherry-CRY-TEVΔ/G79E+T173A+S219V | pRSII413 | Gal | tCYC1 | Yeast / episomally | mCherry | HIS3 selectable marker | Amp | S. Fig 10B |
| p40 | mCherry-TEV/S219V | pRSII413 | Gal | tCYC1 | Yeast / episomally | mCherry | HIS3 selectable marker | Amp | Fig. 3B / S. Fig. 8A |
| p41 | mCherry-TEV/S153N+S219V | pRSII413 | Gal | tCYC1 | Yeast / episomally | mCherry | HIS3 selectable marker | Amp | Fig. 3B / S. Fig. 8A |
| p42 | mCherry-TEV/T30A+S153N+S219V | pRSII413 | Gal | tCYC1 | Yeast / episomally | mCherry | HIS3 selectable marker | Amp | Fig. 3B |
| p43 | mCherry-TEV/G79E+T173A+S219V | pRSII413 | Gal | tCYC1 | Yeast / episomally | mCherry | HIS3 selectable marker | Amp | S. Fig 10A |
| p44 | mtagBFPII-CRY-TEVΔ | pRS-derived | TDH3 | tCYC1 | Yeast / AscI digestion to integrate | mtagBFPII | LEU2 selectable marker | Amp | Fig. 2G / S. Fig. 5B, C |
| p45 | mtagBFPII-CRY-TEVΔ/S219V+S153N | pRS-derived | TDH3 | tCYC1 | Yeast / AscI digestion to integrate | mtagBFPII | LEU2 selectable marker | Amp | Fig. 2G / S. Fig. 5B, C |
| p46 | mtagBFPII-CRY-TEVΔ/s219V+T30A+S153N | pRS-derived | TDH3 | tCYC1 | Yeast / AscI digestion to integrate | mtagBFPII | LEU2 selectable marker | Amp | Fig. 2G / S. Fig. 5B, C |
| p47 | STE2Δ-mCherry-CIBN-PIF6-eLOV-TEVcs-LexAVP16 | pRSII413 | Gal | tCYC1 | Yeast / episomally | mCherry | HIS3 selectable marker | Amp | Fig. 2F |
| p48 | STE2Δ-mCherry-CIBN-PIF6-eLOV-TEVcs-Ala P6-LexAVP16 | pRSII413 | Gal | tCYC1 | Yeast / episomally | mCherry | HIS3 selectable marker | Amp | Fig. 2F |
| p49 | STE2Δ-mCherry-CIBN-PIF6-eLOV-TEVcs-Ala P3-LexAVP16 | pRSII413 | Gal | tCYC1 | Yeast / episomally | mCherry | HIS3 selectable marker | Amp | Fig. 2F |
| p50 | STE2Δ-mCherry-CIBN-PIF6-eLOV-TEVcs-Ala P1-LexAVP16 | pRSII413 | Gal | tCYC1 | Yeast / episomally | mCherry | HIS3 selectable marker | Amp | Fig. 2F |
| p51 | MBP-TEVcs-7xHis-TEVΔ/S219V | pRK793 | tac P | | E. Coli | | | Amp | Fig. 2C, D, E |
| p52 | MBP-TEVcs-7xHis-TEVΔ/S219V+S153N | pRK793 | tac P | | E. Coli | | | Amp | Fig. 2C, D, E |
| p53 | MBP-TEVcs-7xHis-TEVΔ/S219V+T30A+S153N | pRK793 | tac P | | E. Coli | | | Amp | Fig. 2C, D, E |
| p54 | MBP-TEVcs(ENLYFQ/S)-eGFP | pYFJ16 | tac P | | E. Coli | | | Amp | Fig. 3E, G |
| p55 | UAS-mCherry | AAV | UAS | WPRE/PolyA | mammalian / HEK | | | Amp | Fig. 4E, G / S. Fig. 11A, 13 |
| p56 | CD4-HA-CIBN-2xMKII-NNES-heLOV-GAL4 | AAV | CMV | PolyA | mammalian / HEK | HA | | Amp | Fig. 4E, G / S. Fig. 11A, 14 |
| p57 | CMV-eGFP-CaM-TEVΔ/S219V | AAV | CMV | PolyA | mammalian / HEK | V5 / eGFP | | Amp | Fig. 4D / S. Fig. 11A, B. |
| p58 | CMV-eGFP-CaM-TEVΔ/S219V+S153N | AAV | CMV | PolyA | mammalian / HEK | V5 / eGFP | | Amp | Fig. 4D / S. Fig. 11A, B. |
| p59 | CMV-eGFP-CaM-TEVΔ/S219V+T30A+S153N | AAV | CMV | PolyA | mammalian / HEK | V5 / eGFP | | Amp | Fig. 4D / S. Fig. 11A, B. |
| p60 | CMV-eGFP-CaM-TEVΔ/S219V+N177Y | AAV | CMV | PolyA | mammalian / HEK | V5 / eGFP | | Amp | Fig. 4D / S. Fig. 11A, B. |
| p61 | CMV-eGFP-CaM-TEVΔ/S219V+T30A | AAV | CMV | PolyA | mammalian / HEK | V5 / eGFP | | Amp | Fig. 4D / S. Fig. 11A, B. |
| p62 | TRE-mCherry | AAV | TRE | WPRE/PolyA | mammalian / Neuron | | | Amp | Fig. 4F / S. Fig. 12A |
| p63 | tNeu-CIBN-2xMKII-heLOV-tTA | AAV | Syn | PolyA | mammalian / Neuron | HA/ flag/VP16 | | Amp | Fig. 4F / S. Fig. 12A |
| p64 | AAV-EGFP-CaM5f-V5-TEVΔ/S219V | AAV | Syn | WPRE/PolyA | mammalian / Neuron | V5/ eGFP | | Amp | Fig. 4F / S. Fig. 12A |
| p65 | AAV-eGFP-CaM5f-V5-TEVΔ/S219V+S153N | AAV | Syn | WPRE/PolyA | mammalian / Neuron | V5/ eGFP | | Amp | Fig. 4F / S. Fig. 12A |
| p66 | AAV-EGFP-CaM5f-V5-TEVΔ/S219V+T30A+S153N | AAV | Syn | WPRE/PolyA | mammalian / Neuron | V5/ eGFP | | Amp | |
| p67 | AAV-EGFP-CaM5f-V5-TEVΔ/S219V+N177Y | AAV | Syn | WPRE/PolyA | mammalian / Neuron | V5/ eGFP | | Amp | |
| p68 | AAV-EGFP-CaM5f-V5-TEVΔ/S219V+T30A | AAV | Syn | WPRE/PolyA | mammalian / Neuron | V5/ eGFP | | Amp | |
| p69 | B2Ar-heLOV-TEVcs-Gal4-V5 | plx208 | CMV | | mammalian / HEK | HA/ eGFP | Hygromycin gene deleted | Amp | Fig. 4G / S. Fig. 13 |
| p70 | EGFP-p2a-HA-arrestin-TEVΔ/S219V | plx208 | CMV | | mammalian / HEK | HA/ eGFP | Hygromycin gene deleted | Amp | Fig. 4G / S. Fig. 14 |
| p71 | EGFP-p2a-HA-arrestin-TEVΔ/S153N | plx208 | CMV | | mammalian / HEK | HA/ eGFP | Hygromycin gene deleted | Amp | Fig. 4G / S. Fig. 15 |
| p72 | pAAV1 | | | | | | | Amp | Fig. 4F / S. Fig. 12A |
| p73 | pAAV2 | | | | | | | Amp | Fig. 4F / S. Fig. 12A |
| p74 | pDF6 | | | | | | | Amp | Fig. 4F / S. Fig. 12A |
| p75 | mCherry-TEV/S153N+T180A+S219V | pRSII413 | Gal | tCYC1 | Yeast / episomally | mCherry | HIS3 selectable marker | Amp | Fig. 3D / S. Fig. 8A |
| p76 | mCherry-TEV/S135F+S153N+T180A+S219V | pRSII413 | Gal | tCYC1 | Yeast / episomally | mCherry | HIS3 selectable marker | Amp | Fig. 3D / S. Fig. 8A |
| p77 | mCherry-TEV/I138T+S153N+T180A+S219V | pRSII413 | Gal | tCYC1 | Yeast / episomally | mCherry | HIS3 selectable marker | Amp | Fig. 3D / S. Fig. 8A |
| p78 | mCherry-TEV/S135F+I138T+S153N+T180A+S219V | pRSII413 | Gal | tCYC1 | Yeast / episomally | mCherry | HIS3 selectable marker | Amp | Fig. 3D / S. Fig. 8A |
| p79 | mCherry-TEV/I138T+S153N+S219V | pRSII413 | Gal | tCYC1 | Yeast / episomally | mCherry | HIS3 selectable marker | Amp | Fig. 3D / S. Fig. 8A |
| p80 | mtagBFPII-TEV/S153N+T180A+S219V | pRS-derived | TDH3 | tCYC1 | Yeast / AscI digestion to integrate | mtagBFPII | HIS3 selectable marker | Amp | S. Fig. 9 |
| p81 | mtagBFPII-TEV/S135F+S153N+T180A+S219V | pRS-derived | TDH3 | tCYC1 | Yeast / AscI digestion to integrate | mtagBFPII | HIS3 selectable marker | Amp | S. Fig. 9 |
| p82 | mtagBFPII-TEV/I138T+S153N+T180A+S219V | pRS-derived | TDH3 | tCYC1 | Yeast / AscI digestion to integrate | mtagBFPII | HIS3 selectable marker | Amp | S. Fig. 9 |
| p83 | mtagBFPII-TEV/S135F+I138T+S153N+T180A+S219V | pRS-derived | TDH3 | tCYC1 | Yeast / AscI digestion to integrate | mtagBFPII | HIS3 selectable marker | Amp | S. Fig. 9 |
| p84 | mtagBFPII-TEV/I138T+S153N+S219V | pRS-derived | TDH3 | tCYC1 | Yeast / AscI digestion to integrate | mtagBFPII | HIS3 selectable marker | Amp | S. Fig. 9 |
| p85 | mtagBFPII-TEV/I153F+S153N+S219V | pRS-derived | TDH3 | tCYC1 | Yeast / AscI digestion to integrate | mtagBFPII | HIS3 selectable marker | Amp | S. Fig. 9 |
| p86 | STE2Δ-mCherry-CIBN-PIF6-eLOV-TEVcs-Ala P6-LexAVP16 | pRSII413 | Gal | tCYC1 | Yeast / episomally | mCherry | HIS3 selectable marker | Amp | S. Fig. 9 |
| p87 | STE2Δ-mCherry-CIBN-PIF6-eLOV-TEVcs-Ala P3-LexAVP16 | pRSII413 | Gal | tCYC1 | Yeast / episomally | mCherry | HIS3 selectable marker | Amp | S. Fig. 9 |
| p88 | STE2Δ-mCherry-CIBN-PIF6-eLOV-TEVcs-Ala P1-LexAVP16 | pRSII413 | Gal | tCYC1 | Yeast / episomally | mCherry | HIS3 selectable marker | Amp | S. Fig. 9 |
| p89 | STE2Δ-mCherry-CIBN-PIF6-eLOV-TEVcs-LexAVP16 | pRSII413 | Gal | tCYC1 | Yeast / episomally | mCherry | HIS3 selectable marker | Amp | S. Fig. 9 |
| p90 | mCherry-TEV/L56V+S135G | pRSII413 | Gal | tCYC1 | Yeast / episomally | mCherry | HIS3 selectable marker | Amp | S. Fig. 8B |

| ID | Plasmid name | Plasmid Vect | Promoter | Terminator | Expression in | Tags | More Details | Ab. R | Used in figures |
|---|---|---|---|---|---|---|---|---|---|
| p91 | mCherry-TEV/L56V+S135G+S219V | pRSII413 | Gal | tCYC1 | Yeast / episomally | mCherry | HIS3 selectable marker | Amp | S. Fig. 8B |
| p92 | HIS6-MBP-TEV | pYFJ16 | tac P | | E. Coli | MBP / His6 | S219V | Amp | Fig. 4E, G. |
| p93 | HIS6-MBP-TEV I138T S153N T180A | pYFJ16 | tac P | | E. Coli | MBP / His6 | S219V | Amp | Fig. 4E, G. |
| p94 | mCherry-TVMV-tCYC1 | pRSII413 | Gal | | Yeast / episomally | mCherry | HIS3 selectable marker | Amp | S. Fig. 14 |
| p95 | STE2Δ-BFP-CIBN-MKII-eLOV-TVMV(ETVRFQS)-LexAVP16 | pRS-derived | TDH3 | tCYC1 | Yeast / AscI digestion to integrate | BFP | LEU2 selectable marker | Amp | S. Fig. 14 |
| p96 | STE2Δ-BFP-CIBN-MKII-eLOV-TVMV(P1'=M)-LexAVP16 | pRS-derived | TDH3 | tCYC1 | Yeast / AscI digestion to integrate | BFP | LEU2 selectable marker | Amp | S. Fig. 14 |
| p97 | STE2Δ-BFP-CIBN-MKII-eLOV-TVMV(P1'=L)-LexAVP16 | pRS-derived | TDH3 | tCYC1 | Yeast / AscI digestion to integrate | BFP | LEU2 selectable marker | Amp | S. Fig. 14 |
| p98 | STE2Δ-BFP-CIBN-MKII-eLOV-TVMV(P1'=Q)-LexAVP16 | pRS-derived | TDH3 | tCYC1 | Yeast / AscI digestion to integrate | BFP | LEU2 selectable marker | Amp | S. Fig. 14 |
| p99 | LexA-mCherry-VP16 | pRSII413 | Gal | tCYC1 | Yeast / episomally | mCherry | HIS3 selectable marker | Amp | S. Fig. 8B |
| p100 | LexA-mCherry-B42 | pRSII414 | Gal | tCYC1 | Yeast / episomally | mCherry | HIS3 selectable marker | Amp | S. Fig. 8B |
| p101 | LexA-mCherry-Gal4 | pRSII415 | Gal | tCYC1 | Yeast / episomally | mCherry | HIS3 selectable marker | Amp | S. Fig. 8B |
| p102 | MBP-TEVcs(ENLYFQ/M)-eGFP | pYFJ16 | tac P | | E. Coli | MBP/eGFP | | Amp | Fig. 2C, D, E |
| p103 | STE2-citrine | pRS-derived | TDH3 | tCYC1 | Yeast / AscI digestion to integrate | citrine | LEU2 selectable marker | Amp | S. Fig 1A |
| p104 | STE2Δ-citrine | pRS-derived | TDH3 | tCYC1 | Yeast / AscI digestion to integrate | citrine | LEU2 selectable marker | Amp | S. Fig 1A |
| p105 | STE2Δ-BFP-CIBN-eLOV-TEVcs(ENLHFQS, P3=H)-LexAVP16 | pRS-derived | TDH3 | tCYC1 | Yeast / AscI digestion to integrate | BFP / flag | LEU2 selectable marker | Amp | S. Fig 16 |
| p106 | STE2Δ-BFP-CIBN-eLOV-TEVcs(ENLWFQS, P3=W)-LexAVP1 | pRS-derived | TDH3 | tCYC1 | Yeast / AscI digestion to integrate | BFP / flag | LEU2 selectable marker | Amp | S. Fig 16 |
| p107 | STE2Δ-BFP-CIBN-eLOV-TEVcs(ENLYFQS)-LexAVP16 | pRS-derived | TDH3 | tCYC1 | Yeast / AscI digestion to integrate | BFP / flag | LEU2 selectable marker | Amp | Fig. 3B , C / S. Fig. 6C, |
| | | | | | | | | | 7A, B, 8A, B, 10A |
| p108 | mCherry-TEVΔ | AAV | CMV | PolyA | mammalian / HEK | mCherry | | Amp | S. Fig. 5 |
| p109 | mCherry-TEV | AAV | CMV | PolyA | mammalian / HEK | mCherry | | Amp | S. Fig. 5 |
| p110 | mCherry-TEV/S31W+S219V | AAV | CMV | PolyA | mammalian / HEK | mCherry | | Amp | S. Fig. 5 |
| p111 | mCherry-TEV/S153N+S219V | AAV | CMV | PolyA | mammalian / HEK | mCherry | | Amp | S. Fig. 5 |
| p112 | mCherry-TEV/T30A+S153N+S219V | AAV | CMV | PolyA | mammalian / HEK | mCherry | | Amp | S. Fig. 5 |
| p113 | mCherry-TEV-T30I+S219V | AAV | CMV | PolyA | mammalian / HEK | mCherry | | Amp | S. Fig. 5 |
| p114 | mCherry-TEV/N177Y+S219V | AAV | CMV | PolyA | mammalian / HEK | mCherry | | Amp | S. Fig. 5 |
| p115 | CMS817-CMV-mCherry-TEV/T30A S219V | AAV | CMV | PolyA | mammalian / HEK | mCherry | | Amp | S. Fig. 5 |
| p116 | mCherry-TEV/N177Y+T30A+S219V | AAV | CMV | PolyA | mammalian / HEK | mCherry | | Amp | S. Fig. 5 |
| p117 | mCherry-TEV/N177Y+S153N+S219V | AAV | CMV | PolyA | mammalian / HEK | mCherry | | Amp | S. Fig. 5 |
| p118 | mCherry-TEV/N177Y+S31W+S219V | AAV | CMV | PolyA | mammalian / HEK | mCherry | | Amp | S. Fig. 5 |
| p119 | mCherry-TEV/T30A+S153N+N177Y+S219V | AAV | CMV | PolyA | mammalian / HEK | mCherry | | Amp | S. Fig. 5 |
| p120 | mCherry-TEV/T30A+S31W+S219V | AAV | CMV | PolyA | mammalian / HEK | mCherry | | Amp | S. Fig. 5 |
| p121 | mCherry-TEVΔ/S31W+S219V | AAV | CMV | PolyA | mammalian / HEK | mCherry | | Amp | S. Fig. 5 |
| p122 | mCherry-TEVΔ/S153N+S219V | AAV | CMV | PolyA | mammalian / HEK | mCherry | | Amp | S. Fig. 5 |
| p123 | mCherry-TEVΔ/T30A+S153N+S219V | AAV | CMV | PolyA | mammalian / HEK | mCherry | | Amp | S. Fig. 5 |
| p124 | mCherry-TEVΔ/T30I+S219V | AAV | CMV | PolyA | mammalian / HEK | mCherry | | Amp | S. Fig. 5 |
| p125 | mCherry-TEVΔ/N177Y+S219V | AAV | CMV | PolyA | mammalian / HEK | mCherry | | Amp | S. Fig. 5 |
| p126 | mCherry-TEVΔ/T30A+S219V | AAV | CMV | PolyA | mammalian / HEK | mCherry | | Amp | S. Fig. 5 |

### Yeast table

The original strain used is BY4741. **Genotype**: MATa his3Δ1 leu2Δ0 met15Δ0 ura3Δ0. **Description**: S288C-derivative laboratory strain

| ID | Plasmid used | Auxotrophic selection marker |
|---|---|---|
| YS-FRP791 | Addgene plasmid # 58432 (FRP791) | URA3 |
| YS-FRP792 | Addgene plasmid # 58433 (FRP792) | URA3 |
| YS-FRP793 | Addgene plasmid # 58434 (FRP793) | URA3 |
| YS1 | P3-pTDH3-STE2Δ-BFP-CIBN-MKII-eLOV-TEVcs(ENLYFQ/**M**)-LexAVP16-tCYC | URA3/LEU2 |
| YS2 | P105-pTDH3-STE2Δ-BFP-CIBN-MKII-eLOV-TEVcs(ENL**H**FQ/S P3=H)-LexAVP16-tCYC | URA3/LEU2 |
| YS3 | P106-pTDH3-STE2Δ-BFP-CIBN-MKII-eLOV-TEVcs(ENL**W**FQ/S P3=W)-LexAVP16-tCYC | URA3/LEU2 |
| YS4 | P107-pTDH3-STE2Δ-BFP-CIBN-MKII-eLOV-TEVcs(ENLYFQ/**S**)-LexAVP16-tCYC.ape | URA3/LEU2 |
| YS5 | P44-mtagBFPII-CRY-TEVΔ/S219V | URA3/LEU2 |
| YS6 | P45-mtagBFPII-CRY-TEVΔ/S153N+S219V | URA3/LEU2 |
| YS7 | P46-mtagBFPII-CRY-TEVΔ/T30A+S153N+S219V | URA3/LEU2 |
| YS8 | P79-mtagBFPII-TEV/S153N+T180A+S219V | URA3/LEU2 |
| YS9 | P80-mtagBFPII-TEV/S135F+S153N+T180A+S219V | URA3/LEU2 |
| YS10 | P81-mtagBFPII-TEV/I138T+S153N+T180A+S219V | URA3/LEU2 |
| YS11 | P82-mtagBFPII-TEV/S135F+I138T+S153N+T180A+S219V | URA3/LEU2 |
| YS12 | P83-mtagBFPII-TEV/I138T+S153N+S219V | URA3/LEU2 |
| YS13 | P84-mtagBFPII-TEV/I153F+S153N+S219V | URA3/LEU2 |
| YS14 | P94-STE2Δ-BFP-CIBN-MKII-eLOV-TVMVcs(ETVRFQ/S)-LexAVP16 | URA3/LEU2 |
| YS15 | P95-STE2Δ-BFP-CIBN-MKII-eLOV-TVMVcs(ETVRFQ/**M** P1'=M)-LexAVP16 | URA3/LEU2 |
| YS16 | P96-STE2Δ-BFP-CIBN-MKII-eLOV-TVMVcs(ETVRFQ/**L** P1'=L)-LexAVP16 | URA3/LEU2 |
| YS17 | P97-STE2Δ-BFP-CIBN-MKII-eLOV-TVMVcs(ETVRFQ/**Q** P1'=Q)-LexAVP16 | URA3/LEU2 |

| ID | Genetic Background | Used in figures |
|---|---|---|
| YS-FRP791 | BY4741::insul-(lexA-box)1-PminCYC1-Citrine-TCYC1 (URA3) | Fig. 3 |
| YS-FRP792 | BY4741::insul-(lexA-box)2-PminCYC1-Citrine-TCYC1 (URA3) | Fig. 3 |
| YS-FRP793 | BY4741::insul-(lexA-box)4-PminCYC1-Citrine-TCYC1 (URA3) | Fig. 1,2 ,3. |
| YS1 | BY4741::insul-(lexA-box)4-PminCYC1-Citrine-TCYC1 (URA3),pTDH3-STE2Δ-BFP-CIBN-MKII-eLOV-TEVcs(ENLYFQM)-LexAVP16-tCYC (LEU2) | Fig. 1 ,2. / Supp. 1, |
| YS2 | BY4741::insul-(lexA-box)4-PminCYC1-Citrine-TCYC1 (URA3),pTDH3-STE2Δ-BFP-CIBN-MKII-eLOV-TEVcs(ENLHFQS)-LexAVP16-tCYC (LEU2) | Supp. Fig. 15/16 |
| YS3 | BY4741::insul-(lexA-box)4-PminCYC1-Citrine-TCYC1 (URA3),pTDH3-STE2Δ-BFP-CIBN-MKII-eLOV-TEVcs(ENLWFQS)-LexAVP16-tCYC (LEU2) | Supp. Fig. 15/16 |
| YS4 | BY4741::insul-(lexA-box)2-PminCYC1-Citrine-TCYC1 (URA3),pTDH3-STE2Δ-BFP-CIBN-MKII-eLOV-TEVcs(ENLYFQS)-LexAVP16-tCYC (LEU2) | Fig. 3. / Supp. 7, 8, |
| YS5 | BY4741::insul-(lexA-box)4-PminCYC1-Citrine-TCYC1 (URA3), pTDH3-mtagBFPII-TEVΔ/S219V (LEU2) | Supp. Fig. 5C |
| YS6 | BY4741::insul-(lexA-box)4-PminCYC1-Citrine-TCYC1 (URA3), pTDH3-mtagBFPII-TEVΔ/S153N+S219V (LEU2) | Supp. Fig. 5C |
| YS7 | BY4741::insul-(lexA-box)4-PminCYC1-Citrine-TCYC1 (URA3), pTDH3-mtagBFPII-TEVΔ/T30A+S153N+S219V (LEU2) | Supp. Fig. 5C |
| YS8 | BY4741::insul-(lexA-box)4-PminCYC1-Citrine-TCYC1 (URA3), pTDH3-mtagBFPII-TEVΔ/S153N+T180A+S219V (LEU2) | Supp. Fig. 9 |
| YS9 | BY4741::insul-(lexA-box)4-PminCYC1-Citrine-TCYC1 (URA3), pTDH3-mtagBFPII-TEVΔ/S135F+S153N+T180A+S219V (LEU2) | Supp. Fig. 9 |
| YS10 | BY4741::insul-(lexA-box)4-PminCYC1-Citrine-TCYC1 (URA3), pTDH3-mtagBFPII-TEVΔ/I138T+S153N+T180A+S219V (LEU2) | Supp. Fig. 9 |
| YS11 | BY4741::insul-(lexA-box)4-PminCYC1-Citrine-TCYC1 (URA3), pTDH3-mtagBFPII-TEVΔ/S135F+I138T+S153N+T180A+S219V (LEU2) | Supp. Fig. 9 |
| YS12 | BY4741::insul-(lexA-box)4-PminCYC1-Citrine-TCYC1 (URA3), pTDH3-mtagBFPII-TEVΔ/I138T+S153N+S219V (LEU2) | Supp. Fig. 9 |
| YS13 | BY4741::insul-(lexA-box)4-PminCYC1-Citrine-TCYC1 (URA3), pTDH3-mtagBFPII-TEVΔ/I153F+S153N+S219V (LEU2) | Supp. Fig. 9 |
| YS14 | BY4741::insul-(lexA-box)4-PminCYC1-Citrine-TCYC1 (URA3),pTDH3-STE2Δ-BFP-CIBN-MKII-eLOV-TEVcs(ETVRFQ/S)-LexAVP16-tCYC (LEU2) | Supp. Fig. 14 |
| YS15 | BY4741::insul-(lexA-box)4-PminCYC1-Citrine-TCYC1 (URA3),pTDH3-STE2Δ-BFP-CIBN-MKII-eLOV-TEVcs(ETVRFQ/M P1'=M)-LexAVP16-tCYC (LEU2) | Supp. Fig. 14 |
| YS16 | BY4741::insul-(lexA-box)4-PminCYC1-Citrine-TCYC1 (URA3),pTDH3-STE2Δ-BFP-CIBN-MKII-eLOV-TEVcs(ETVRFQ/L P1'=L)-LexAVP16-tCYC (LEU2) | Supp. Fig. 14 |
| YS17 | BY4741::insul-(lexA-box)4-PminCYC1-Citrine-TCYC1 (URA3),pTDH3-STE2Δ-BFP-CIBN-MKII-eLOV-TEVcs(ETVRFQ/Q P1'=Q)-LexAVP16-tCYC (LEU2) | Supp. Fig. 14 |

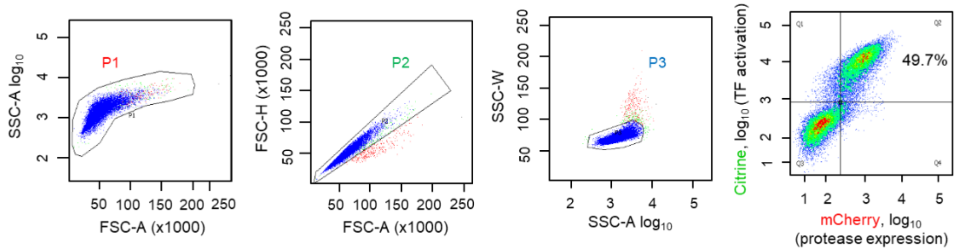


Tube: Example

| Population | #Events | %Parent | %Total |
|---|---|---|---|
| All Events | 20,000 | #### | 100.0 |
| P1 | 19,995 | 100.0 | 100.0 |
| P2 | 19,709 | 98.6 | 98.5 |
| P3 | 19,554 | 99.2 | 97.8 |
| Q1 | 1,319 | 6.6 | 6.6 |
| Q2 | 8,616 | 43.1 | 43.1 |
| Q3 | 9,389 | 46.9 | 46.9 |
| Q4 | 676 | 3.4 | 3.4 |

**Sample FACS plots showing the gating parameters**. Percentage values reflect the fraction of cells with high Citrine intensity, i.e., cells in the upper FACS quadrants Q1 + Q2.

**TEV sequences**

TEVΔ-S219V

Ggagaatccttgtttaagggaccacgtgattacaacccgatatcgagcaccatttgtcatttgacgaatgaatctgatgggcacacaacatcgttgtatggtattggattt
ggtcccttcatcattacaaacaagcacttgtttagaagaaataatggaacactgttggtccaatcactacatggtgtattcaaggtcaagaacaccacgactttgcaaca
acacctcattgatgggagggacatgataattattcgcatgcctaaggatttcccaccatttcctcaaaagctgaaatttagagagccacaaagggaagagcgcatatg
tcttgtgacaaccaacttccaaactaagagcatgtctagcatggtgtcagacactagttgcacattcccttcatctgatggcatattctggaagcattggattcaaaccaa
ggatgggcagtgtggcagtccattagtatcaactagagatgggttcattgttggtatacactcagcatcgaatttcaccaacacaaacaattatttcacaagcgtgccga
aaaacttcatggaattgttgacaaatcaggaggcgcagcagtgggttagtggttggcgattaaatgctgactcagtattgtgggggggccataaagttttcatg**gtg.**

GESLFKGPRDYNPISSTICHLTNESDGHTTSLYGIGFGPFIITNKHLFRRNNGTLLVQSLHGVFKVKNTTTLQQHLIDGR
DMIIIRMPKDFPPFPQKLKFREPQREERICLVTTNFQTKSMSSMVSDTSCTFPSSDGIFWKHWIQTKDGQCGSPLVST
RDGFIVGIHSASNFTNTNNYFTSVPKNFMELLTNQEAQQWVSGWRLNADSVLWGGHKVFM**V**

TEV1Δ-(S135N/S219V)

Ggagaatccttgtttaagggaccacgtgattacaacccgatatcgagcaccatttgtcatttgacgaatgaatctgatgggcacacaacatcgttgtatggtattggattt
ggtcccttcatcattacaaacaagcacttgtttagaagaaataatggaacactgttggtccaatcactacatggtgtattcaaggtcaagaacaccacgactttgcaaca
acacctcattgatgggagggacatgataattattcgcatgcctaaggatttcccaccatttcctcaaaagctgaaatttagagagccacaaagggaagagcgcatatg
tcttgtgacaaccaacttccaaactaagagcatgtctagcatggtgtcagacactagttgcacattcccttcatctgatggcatattctggaagcattggattcaaaccaa
ggatgggcagtgtggc**aat**ccattagtatcaactagagatgggttcattgttggtatacactcagcatcgaatttcaccaacacaaacaattatttcacaagcgtgccga
aaaacttcatggaattgttgacaaatcaggaggcgcagcagtgggttagtggttggcgattaaatgctgactcagtattgtgggggggccataaagttttcatg**gtg.**

GESLFKGPRDYNPISSTICHLTNESDGHTTSLYGIGFGPFIITNKHLFRRNNGTLLVQSLHGVFKVKNTTTLQQHLIDGR
DMIIIRMPKDFPPFPQKLKFREPQREERICLVTTNFQTKSMSSMVSDTSCTFPSSDGIFWKHWIQTKDGQCG**N**PLVST
RDGFIVGIHSASNFTNTNNYFTSVPKNFMELLTNQEAQQWVSGWRLNADSVLWGGHKVFM**V**

TEV2Δ-(T30A/S135N/S219V)

Ggagaatccttgtttaagggaccacgtgattacaacccgatatcgagcaccatttgtcatttgacgaatgaatctgatgggcacaca**gca**tcgttgtatggtattggattt
ggtcccttcatcattacaaacaagcacttgtttagaagaaataatggaacactgttggtccaatcactacatggtgtattcaaggtcaagaacaccacgactttgcaaca
acacctcattgatgggagggacatgataattattcgcatgcctaaggatttcccaccatttcctcaaaagctgaaatttagagagccacaaagggaagagcgcatatg
tcttgtgacaaccaacttccaaactaagagcatgtctagcatggtgtcagacactagttgcacattcccttcatctgatggcatattctggaagcattggattcaaaccaa
ggatgggcagtgtggcaatccattagtatcaactagagatgggttcattgttggtatacactcagcatcgaatttcaccaacacaaacaattatttcacaagcgtgccga
aaaacttcatggaattgttgacaaatcaggaggcgcagcagtgggttagtggttggcgattaaatgctgactcagtattgtgggggggccataaagttttcatg**gtg.**

GESLFKGPRDYNPISSTICHLTNESDGHT**A**SLYGIGFGPFIITNKHLFRRNNGTLLVQSLHGVFKVKNTTTLQQHLIDGR
DMIIIRMPKDFPPFPQKLKFREPQREERICLVTTNFQTKSMSSMVSDTSCTFPSSDGIFWKHWIQTKDGQCG**N**PLVST
RDGFIVGIHSASNFTNTNNYFTSVPKNFMELLTNQEAQQWVSGWRLNADSVLWGGHKVFM**V**

TEV-S219V

Ggagaatccttgtttaagggaccacgtgattacaacccgatatcgagcaccatttgtcatttgacgaatgaatctgatgggcacacaacatcgttgtatggtattggattt
ggtcccttcatcattacaaacaagcacttgtttagaagaaataatggaacactgttggtccaatcactacatggtgtattcaaggtcaagaacaccacgactttgcaaca
acacctcattgatgggagggacatgataattattcgcatgcctaaggatttcccaccatttcctcaaaagctgaaatttagagagccacaaagggaagagcgcatatg
tcttgtgacaaccaacttccaaactaagagcatgtctagcatggtgtcagacactagttgcacattcccttcatctgatggcatattctggaagcattggattcaaaccaa
ggatgggcagtgtggcagtccattagtatcaactagagatgggttcattgttggtatacactcagcatcgaatttcaccaacacaaacaattatttcacaagcgtgccga
aaaacttcatggaattgttgacaaatcaggaggcgcagcagtgggttagtggttggcgattaaatgctgactcagtattgtgggggggccataaagttttcatg**gtt**aaa
cctgaagagccttttcagccagttaaggaagcgactcaactcatgaatgaattggtctacagccag.

GESLFKGPRDYNPISSTICHLTNESDGHTTSLYGIGFGPFIITNKHLFRRNNGTLLVQSLHGVFKVKNTTTLQQHLIDGR
DMIIIRMPKDFPPFPQKLKFREPQREERICLVTTNFQTKSMSSMVSDTSCTFPSSDGIFWKHWIQTKDGQCGSPLVST
RDGFIVGIHSASNFTNTNNYFTSVPKNFMELLTNQEAQQWVSGWRLNADSVLWGGHKVFM**V**KPEEPFQPVKEATQL
MNELVYSQ

uTEV3-(I138T/S153N/T180A/S219V)

ggagaatccttgtttaagggaccacgtgattacaacccgatatcgagcaccatttgtcatttgacgaatgaatctgatgggcacacaacatcgttgtatggtattggatttg
gtcccttcatcattacaaacaagcacttgtttagaagaaataatggaacactgttggtccaatcactacatggtgtattcaaggtcaagaacaccacgactttgcaacaa
cacctcattgatgggagggacatgataattattcgcatgcctaaggatttcccaccatttcctcaaaagctgaaatttagagagccacaaagggaagagcgcatatgt
cttgtgacaaccaacttccaaactaagagcatgtctagcatggtgtcagacactagttgcacattcccttcatctgatggc**aca**ttctggaagcattggattcaaaccaa
ggatgggcagtgtggc**aat**ccattagtatcaactagagatgggttcattgttggtatacactcagcatcgaatttcaccaacacaaacaattatttc**gca**agcgtgccga
aaaacttcatggaattgttgacaaatcaggaggcgcagcagtgggttagtggttggcgattaaatgctgactcagtattgtgggggggccataaagttttcatg**gtg**aa
acctgaagagccttttcagccagttaaggaagcgactcaactcatgaatgaattggtctacagccag

GESLFKGPRDYNPISSTICHLTNESDGHTTSLYGIGFGPFIITNKHLFRRNNGTLLVQSLHGVFKVKNTTTLQQHLIDGR
DMIIIRMPKDFPPFPQKLKFREPQREERICLVTTNFQTKSMSSMVSDTSCTFPSSDG**T**FWKHWIQTKDGQCG**N**PLVST
RDGFIVGIHSASNFTNTNNYF**A**SVPKNFMELLTNQEAQQWVSGWRLNADSVLWGGHKVFM**V**KPEEPFQPVKEATQL
MNELVYSQ
